# Individual listening success explained by synergistic interaction of two distinct neural filters

**DOI:** 10.1101/512251

**Authors:** Sarah Tune, Lorenz Fiedler, Mohsen Alavash, Jonas Obleser

## Abstract

Successful speech comprehension requires the listener to differentiate relevant from irrelevant sounds. Recent neurophysiological studies have typically addressed one of two candidate neural filter solutions for this problem: the selective neural tracking of speech in auditory cortex via the modulation of phase-locked cortical responses, or the suppression of irrelevant inputs via alpha power modulations in parieto-occipital cortex. However, empirical evidence on their relationship and direct relevance to behavior is scarce. Here, a large, age-varying sample (N=76, 39–70 years) underwent a challenging dichotic listening task. Irrespective of listeners’ age, measures of behavioral performance, neural speech tracking, and alpha power lateralization all increased in response to spatial-attention cues. Under most challenging conditions, individual listening success was predicted best by the synergistic interaction of these two distinct neural filter strategies. Trial-by-trial fluctuations of lateralized alpha power and ignored-speech tracking did not co-vary, which demonstrates two neurobiologically distinct filter mechanisms.

## Introduction

Real-life listening is characterized by the concurrence of sound sources that compete for our attention (Cherry, 1953). Successful speech comprehension thus relies on the selective filtering of relevant and irrelevant inputs. How does the brain instantiate attentional filter mechanisms that allow for the controlled inhibition and amplification of speech? Recent neuroscientific approaches to this question have focused on two different neural filter strategies originating from distinct research traditions:

From the visual domain stems an influential line of research that suggests a domain-general role of parieto-occipital alpha-band (~8–12 Hz) oscillatory activity in implementing selective attention. Here, the key assumption is that increases in alpha power support the controlled, top-down suppression of behaviorally-irrelevant information (Rihs et al., 2007; Jensen and Mazaheri, 2010; Foxe and Snyder, 2011; Händel et al., 2011). Accordingly, listening tasks that require spatially-directed attention are neurally supported by a hemispheric lateralization of changes in alpha power over occipital, parietal but also sensory cortices (Kerlin et al., 2010; Müller and Weisz, 2011; Ahveninen et al., 2013; Wöstmann et al., 2016; Tune et al., 2018; Wöstmann et al., 2018).

In addition, there is a prominent line of research that focuses on the role of low-frequency (1–8 Hz) neural activity in and around auditory cortex in the selective representation of sound inputs. It is assumed that slow cortical dynamics temporally align with (or “neurally track”) auditory input signals to allow for optimal processing of behaviorally-relevant sensory information (Schroeder and Lakatos, 2009; Schroeder et al., 2010; Henry and Obleser, 2012). In human speech comprehension, recent studies found evidence for the preferential neural tracking of attended compared to ignored speech in superior temporal brain areas close to auditory cortex (Ding and Simon, 2012; Mesgarani and Chang, 2012; Horton et al., 2013; Zion Golumbic et al., 2013; O’Sullivan et al., 2014).

However, with few exceptions, these two proposed neural filter strategies have been studied independently of one another (but see Kerlin et al., 2010). Also, they have often been studied using tasks that are difficult to relate to natural listening situations (Lakatos et al., 2016; Henry et al., 2017). We thus lack understanding whether or how modulations in (lateralized) alpha power, presumably arising from domain-general networks involving parietal cortex, and the neural tracking of attended and ignored speech in wider auditory cortex interact in the service of successful listening behavior. At the same time, few studies using more real-life listening and speech-tracking measures were able to explicitly address the functional relevance of the discussed neural filter strategies, that is, their immediate consequences for speech comprehension success (but see Mesgarani and Chang, 2012).

In the present EEG study, we aim at closing these gaps by leveraging the statistical power and representativeness of a large, age-varying participant sample. We use a novel dichotic listening paradigm to enable a synoptic look at concurrent changes in alpha power and neural speech tracking at the single-trial level. More specifically, our linguistic variant of a classic Posner paradigm (Posner, 1980) emulates a challenging dual-talker listening situation in which speech comprehension is supported by two different listening cues (Alavash et al., 2018). These cues encourage the use of two complementary cognitive strategies to improve comprehension: A spatial-attention cue guides auditory attention in space, whereas a semantic cue affords more specific semantic predictions of upcoming speech. Previous research has shown that the sensory analysis of speech and, to a lesser degree, the modulation of alpha power are influenced by the availability of higher-order linguistic information (Obleser and Weisz, 2012; Sohoglu et al., 2012; Peelle et al., 2013; Presacco et al., 2016a; Wöstmann et al., 2017).

Both cues were presented in an informative and uninformative version, allowing us to examine relative changes in listening success and in the modulation of neural measures thought to enable auditory selective attention. Based on the neural and behavioral results, we focused on four research questions (see Fig. 1).

**Figure 1.**
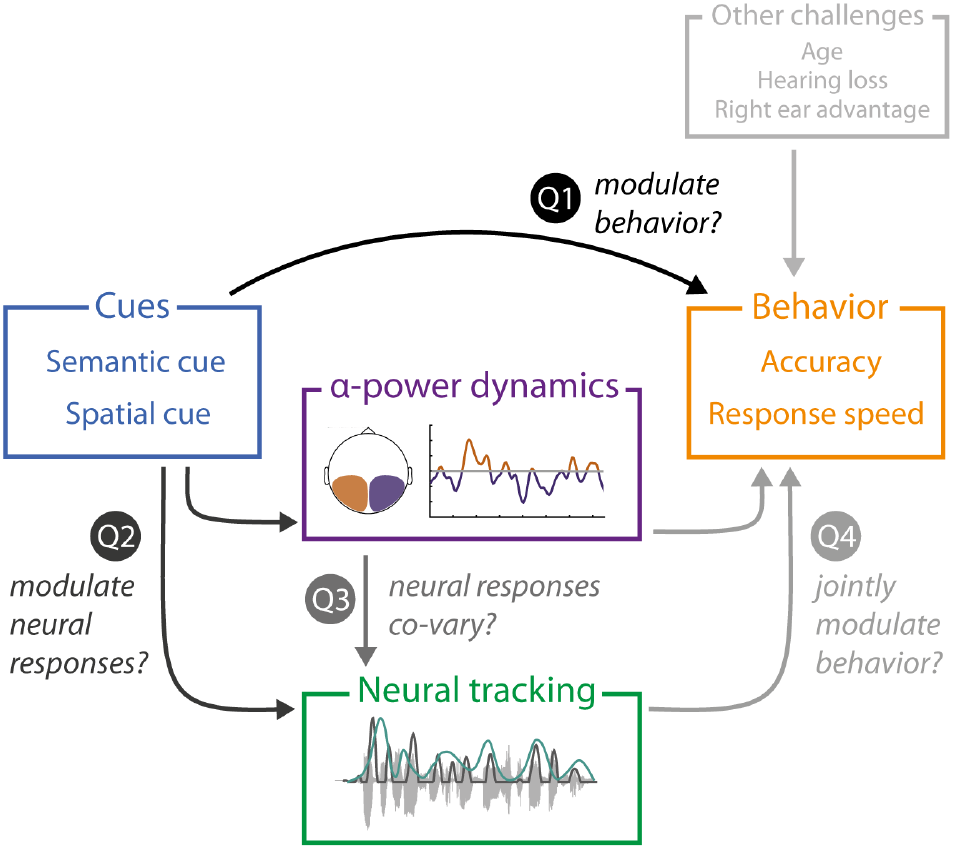
Schematic illustration of the research questions addressed in the present study. The dichotic listening task manipulated the attentional focus and semantic predictability of upcoming input using two separate visual cues. We investigated whether informative cues would enhance behavioral performance (Q1). In line with (Q2), we also examined the degree to which listening cues modulated the two neural measures of interest: neural tracking and lateralization of alpha power. Finally, we assessed (Q3) the co-variation of neural measures, and (Q4) their potency in predicting behavioral performance. Furthermore, we controlled for additional factors that may challenge listening success and its underlying neural strategies.

First, we predicted that informative listening cues should increase speech comprehension success: These cues allow to deploy auditory selective attention (compared to divided attention), and to generate more specific (compared to only general) semantic predictions, respectively.

Secondly, we asked how the different cue-cue combinations would modulate the two key neural measures—alpha power lateralization and neural speech tracking. We hypothesized that selective (compared to divided) attention should increase the strength of alpha power lateralization and neural tracking of the to-be-attended speech signal, respectively.

An important and often neglected third research question pertains to a direct, trial-by-trial relationship of these two candidate neural measures: Do changes in alpha power lateralization impact the degree to which attended and ignored speech signals are neurally tracked by low-frequency cortical responses?

Our most important final research question asked whether the observed (co-)variation of alpha power and neural speech tracking of attended and ignored speech would in turn allow us to predict single-trial behavioral success in this challenging listening situation. In addressing these research questions, we acknowledge additional, potentially detrimental influences on listening success and its supporting neural strategies. These include age, hearing loss, or hemispheric asymmetries in speech processing due to the well-known right-ear advantage (Kimura, 1961; Broadbent and Gregory, 1964).

The following figure supplements are available for figure 2:

**Figure 2.**
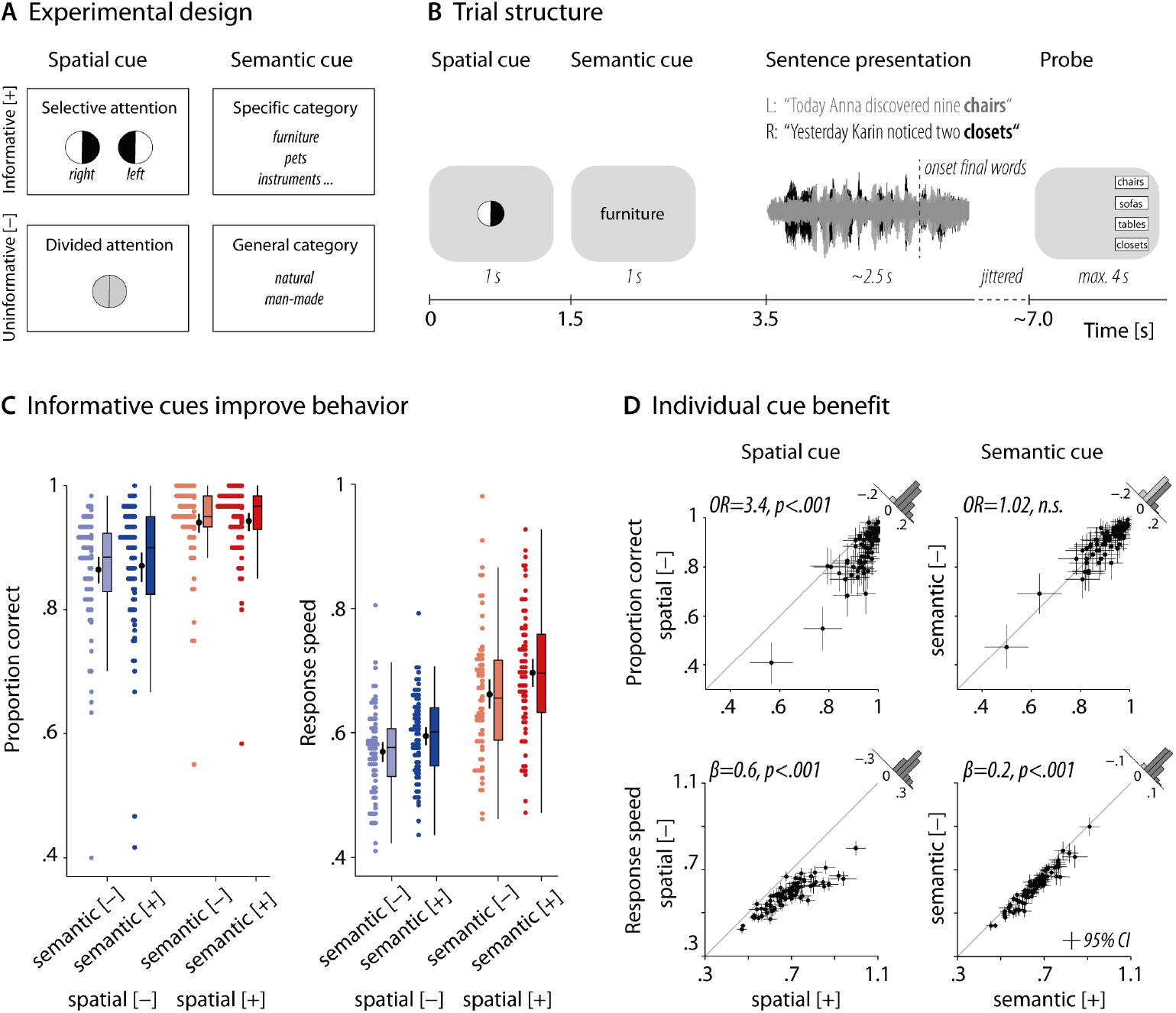
Experimental design and behavioral benefit from informative cues. (A) Visualization of the employed 2×2 design. The level of the spatial and semantic cue differed on a trial-by-trial basis. Top row shows the informative [+] cue levels, bottom row the uninformative [-] cue levels. (B) Schematic representation of the temporal order of events for a given trial. Successive display of the two visual cues precedes the dichotic presentation of two sentences uttered by the same female talker. After sentence presentation, participants had to select the final word from four alternative words. (C) Grand average and individual results of accuracy and response speed shown per cue-cue combination. Colored dots are individual (N=76) trial-averages, black dots and vertical lines show individual means ± bootstrapped 95% confidence intervals. (D) Individual cue benefits for all 76 participants displayed separately for the two cues and dependent behavioral measures, respectively. Across participants, we observed a consistent cue benefit of the informative compared to the uninformative spatial-attention cue on both accuracy and response speed. Black dots indicate individual trial-averages ± bootstrapped 95% confidence intervals. Histograms show the distribution of the difference for informative vs. uninformative levels. OR: Odds ratio parameter estimate from generalized linear mixed-effects models; *β*: slope parameter estimate from general linear mixed-effects models. The following figure supplements are available for figure 2:

**Figure supplement 1.** Histogram showing age distribution of N=76 participants across six age bins.
**Figure supplement 2.** Individual and mean air-conduction thresholds (PTA) averaged across the left and right ear.
**Figure supplement 3.** Right-ear advantage for accuracy and response speed.

## Results

We recorded EEG from an age-varying sample (N=76) of healthy middle-aged and older adults (39–70 years of age) who performed a challenging dichotic listening task (Alavash et al., 2018). In this linguistic variant of a classic Posner paradigm, participants listened to two competing sentences spoken by the same female talker, and were asked to identify the final word in one of the two sentences. Importantly, sentence presentation was preceded by two visual cues. First, a spatial-attention cue encouraged the use of either selective or divided attention by providing informative or uninformative instructions about the to-be-attended, and thus later probed, ear. The second cue indicated the semantic category that applied to both final target words. The provided category could represent a general or specific level, thus allowing for more or less precise prediction of the upcoming speech signal (Fig. 2A, B).

Our analyses aim at explaining behavioral task performance in a challenging listening situation, and the degree to which it is leveraged by two key neural measures of auditory attention: the lateralization of 8–12 Hz alpha power, and the neural tracking of attended and ignored speech by slow cortical responses. Using generalized linear mixed-effects models on single-trial data, we investigated the cue-driven modulation of behavior and neural measures, as well as the interaction of neural measures, and their (joint) influence on listening success.

### Informative spatial cue improves listening success

The analysis of behavioral performance tested the effect of informative versus uninformative listening cues on speech comprehension success. Overall, participants achieved a mean accuracy of 90.5% (sd 7.6%) but as shown in Fig. 2C, the behavioral results varied between the different combination of listening cues. We investigated these relative changes in accuracy and response speed at the single-trial level. As highlighted above, our models consider the potential influence of additional factors such as the probed ear (left vs. right), age, or hearing acuity as measured by the pure-tone average across both ears (PTA; see also Fig. 2–supplements 1 and 2). Variability between participants (see Fig. 2D) and items was accounted for via the inclusion of participant- and item-specific random effects (see Tables S1 and S2 for full model details).

As expected, the analyses revealed a strong behavioral benefit of informative compared to uninformative spatial-attention cues. In selective-attention trials, participants responded more accurately and faster (accuracy: *odds ratio* (*OR*)=*3.37, p*<.001; response speed: *β*=.58, *p*<.001). That is, when cued to one of the two sides, participants responded on average 250 ms faster and their probability of giving a correct answer increased by 5%.

Participants also responded generally faster in trials in which they were given a specific and thus informative semantic cue (*β*=.19, *p*<.001), most likely reflecting a semantic priming effect that led to faster word recognition. A corresponding semantic-cue effect on accuracy was only found for trials in which the onset of the later-probed sentence preceded that of the unprobed sentence (interaction semantic cue × first onset: *OR*=2.29, *p*<.001; simple effect semantic cue (probed-first trials): *OR*=1.59, *p*=.03). Note that these slight time delays between dichotically presented sentences resulted from these sentences being naturally recorded but subsequently onset-aligned for the task-relevant, sentence-final words (see Supplemental Information for details).

As in a previous fMRI implementation of this task (Alavash et al., 2018), we did not find evidence for any interactive effects of the two listening cues on either accuracy or response speed. The inclusion of the respective interaction terms either did not significantly improve model fit (likelihood ratio test; accuracy: *X*^2^_1_=.04; *p*=.84) or did not reach significance in the final best-fitting model (response speed: *β*=.10, *p*=.13).

**Figure 3.**
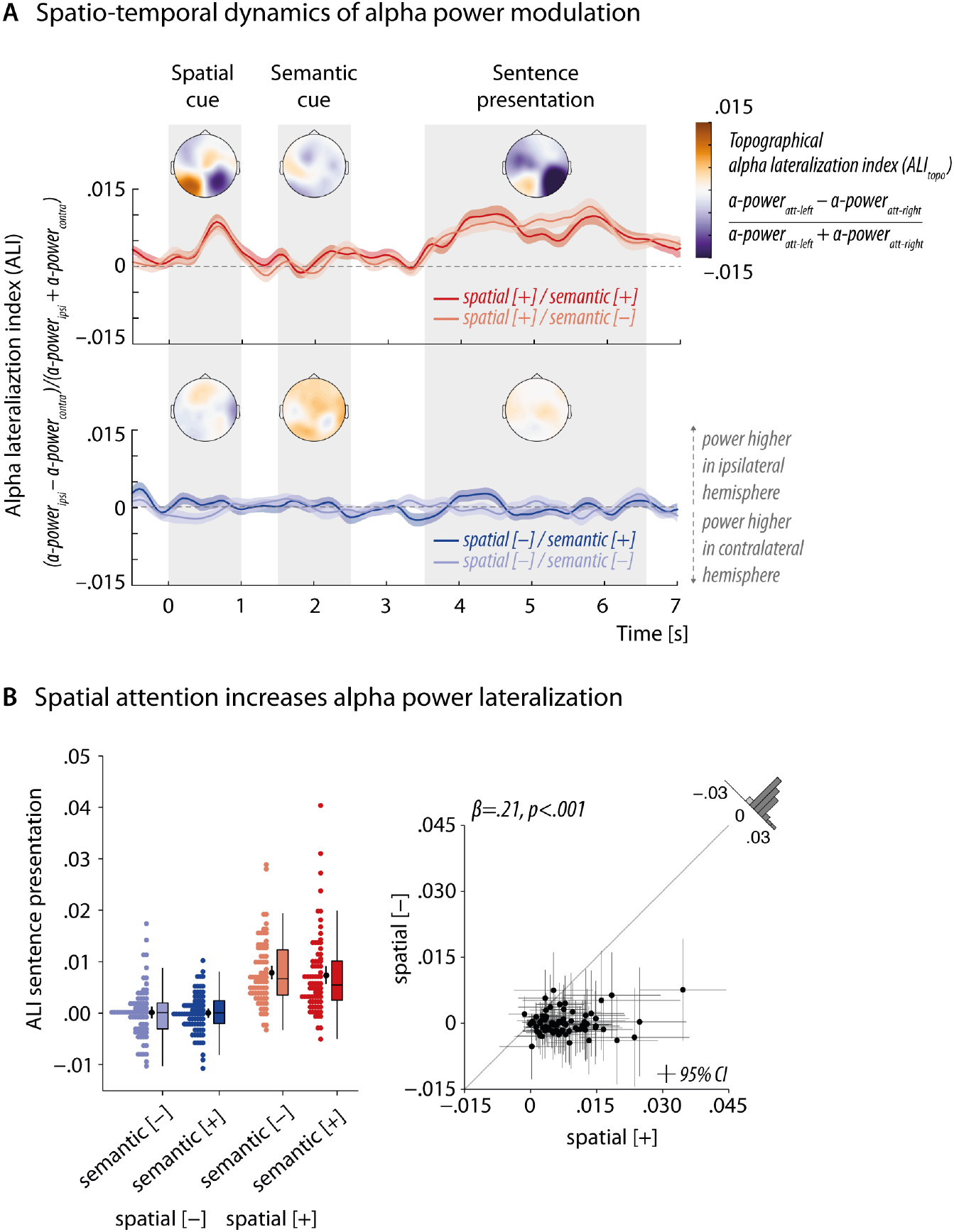
Informative spatial cue elicits increased alpha power lateralization before and during speech presentation. (A) Attentional modulation of 8–12 Hz alpha power shown throughout the trial across all N=76 participants. Topographical maps indicate the spatial distribution of the alpha lateralization index (ALI_topo_) collapsed across the semantic-cue levels in three time windows of interest (spatial cue: 0–1 s; semantic cue: 1.5–2.5 s; sentence presentation: 3.5–6.5 s; shaded in grey). Warm colors indicate higher power in attend-/probed-left compared to attend-/probed-right trials. The opposite relationship holds for cold colors. Red and blue traces show the temporally resolved alpha lateralization index (ALI) for each cue-cue combination. Positive values indicate relatively higher alpha power in the hemisphere ipsilateral to the attended/-probed sentence compared to the contralateral hemisphere. (B) Strength of the ALI during sentence presentation (3.5–6.5 s) shown per cue-cue combination (*right plot*). 1qColored dots represent trial-averaged individual results, black dots and error bars indicate the grand-average and bootstrapped 95% confidence intervals. Consistent increase in alpha power lateralization across all participants (*left plot*). Black dots represent individual mean ALI values ± bootstrapped 95% confidence intervals. Histogram shows distribution of differences in ALI for informative vs. uninformative spatial-cue levels. *β*: slope parameter estimate from the corresponding general linear mixed-effects model. The following figure supplements are available for figure 3:

**Figure supplement 1.** Whole-trial overall oscillatory power averaged across all trials, electrodes, and N = 76 participants.
**Figure supplement 2.** Grand-average whole-trial alpha lateralization index (ALI) calculated without adjustment for overall differences in power across hemispheres.
**Figure supplement 3.** Grand-average whole-trial alpha lateralization index (ALI) time-locked to the onset of the sentence-final target words.

### Lateralization of alpha power modulated by selective attention but not semantic predictability

In line with our second research question, we tested the hypothesis that an informative spatial cue would lead to a lateralization of alpha power in parietal-occipital brain areas. We relied on two established neural indices (Haegens et al., 2011; Mesgarani and Chang, 2012): The topographical alpha lateralization index [ALI_topo_= (α-power_att-left_ – α-power_att-right_) / (α-power_att-left_ + α-power_att-right_)] contrasts alpha power in attend-left vs. attend-right trials (for divided-attention conditions: probed-left vs. probed-right trials) and yields a topographical representation of attention-related changes in power. It is complemented by the alpha lateralization index [ALI = (α-power_ipsi_ – α-power_contra_) / (α-power_ipsi_ + α – power_contra_)] that compares alpha power ipsi- and contralateral to the attended/probed ear to provide a temporally-resolved single-trial measure of alpha power lateralization (for overall whole-trial changes in power, see Fig. 3–supplement 1).

In accordance with the proposed role of alpha power modulation in supporting selective attention, the instruction to pay attention to one of the two sides elicited a pronounced lateralization of 8–12 Hz alpha power over posterior channels during the spatial cue and during the actual sentence presentation (see ALI_topo_ maps in Fig. 3A). The time course of alpha power lateralization per cue-cue combination further supported this observation: We found a strong but transient increase in lateralization in response to an informative spatial-attention cue. After a subsequent break-down of the lateralization during semantic-cue presentation, it re-emerged in time for the start of dichotic sentence presentation and peaked again during the presentation of the task-relevant sentence-final words (see Fig. 3–supplement 3 for ALI time course time-locked to the presentation of sentence-final words).

To investigate the influence of the spatial-attention but also the semantic cue on the degree of alpha power lateralization during sentence presentation in more detail, we extracted single-trial average values from this time window and submitted them as dependent measure to linear mixed-effects analysis (see Fig. 3B and Table S3 for full model details). As expected, the analysis revealed a highly significant and consistent increase in alpha power lateralization for informative compared to uninformative spatial cues (*β*=.21, *p*<.001). Notably, we did not find any evidence of a modulation of alpha lateralization during sentence presentation by the semantic cue nor any joint influence of the spatial and semantic cue (likelihood ratio tests; main effect semantic cue: *X*^2^_1_=32, *p*=.57; interaction of cues: *X*^2^_1_=.19, *p*=.67). Note that the analysis of alpha power lateralization during spatial-cue presentation yielded a comparable increase in lateralization in response to an informative spatial cue (*β*=.08, *p*<.001; see Table S4 for full details).

### Selective tracking of attended and ignored speech

In close correspondence with our analysis of alpha power lateralization, we investigated whether changes in attentional demand and semantic predictability would modulate the neural tracking of attended and ignored speech. To this end, we used linear backward (‘decoding’) models to reconstruct the onset envelopes of the to-be-attended and -ignored sentences (for simplicity hereafter referred to as *attended* and *ignored*). We compared the reconstructed envelopes to the envelopes of presented sentences using Pearson’s correlation. The resulting correlation coefficients are a measure of reconstruction accuracy and serve as an indicator of how strongly the envelopes of attended and ignored sentences were tracked by slow cortical responses (i.e., neural tracking; see Fig. 4A and figure supplement 1). Reconstruction models were trained on selective-attention trials, only, but then utilized to reconstruct attended (probed) and ignored (unprobed) envelopes for both attention conditions (see Methods for details).

**Figure 4.**
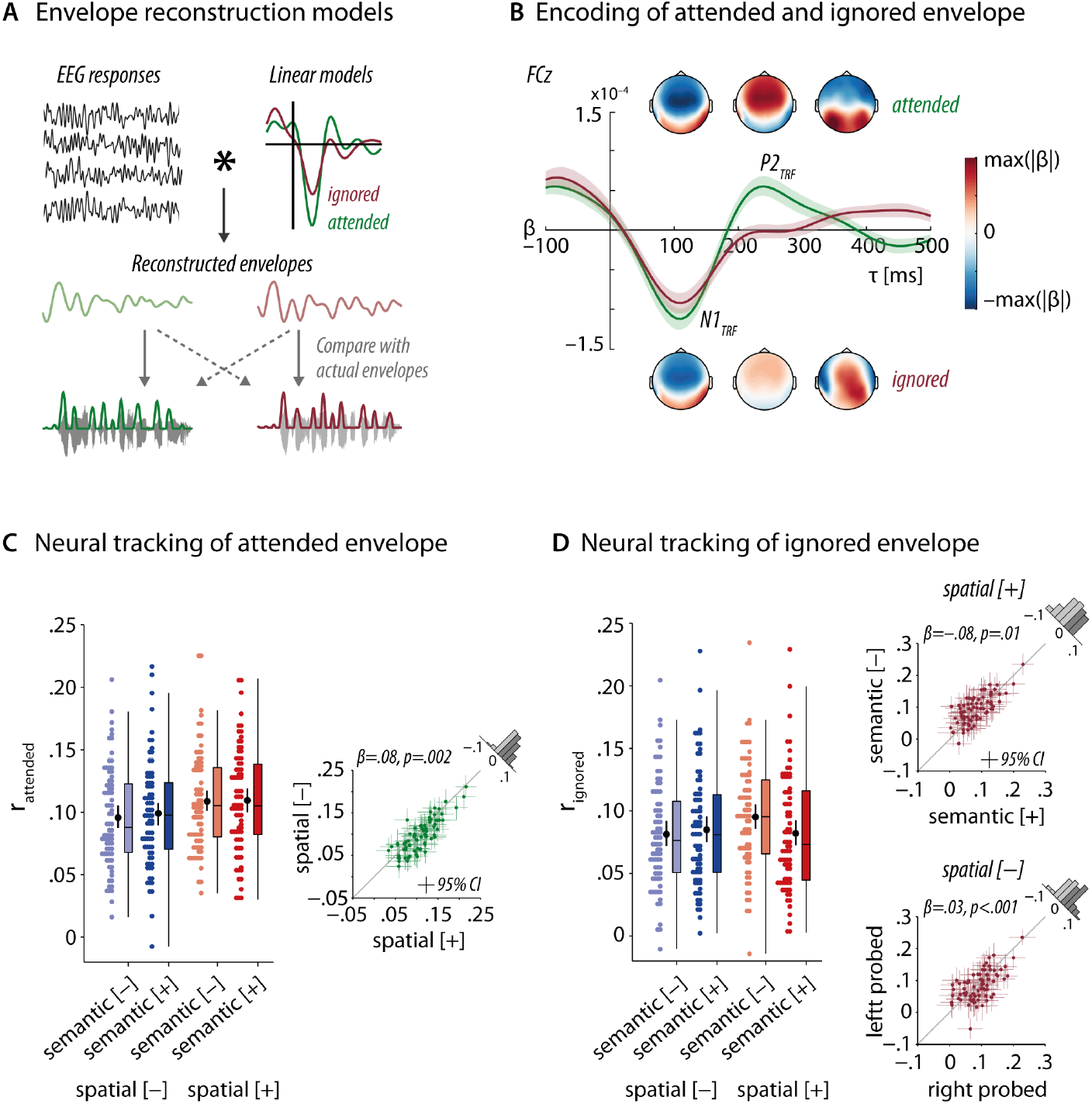
Neural speech tracking of attended and ignored sentences. (A) Schematic representation of the linear backward model approach used to reconstruct onset envelopes on the basis of recorded EEG responses. Following the estimation of linear backward models (see Fig. 4–supplement 1 for details), envelopes of attended and ignored sentences were reconstructed via convolution of EEG responses with the estimated linear backward models at all EEG channels. Reconstructed envelopes were compared to the envelopes of presented sentences to assess reconstruction accuracy. (B) Topographical distribution of linear backward model weights shown in line with forward-transformed temporal response functions for attended (green, top row) and ignored (red, bottom row) speech at electrode FCz. Models are averaged across all N=76 participants, and across attend-left and attend-right trials. Topographical maps correspond to the time lag intervals of 90–110 ms, 200–250 ms, and 350–450 ms. Error bands indicate 95% confidence intervals. (C, D) Changes in neural tracking of attended and ignored speech across cue-cue combinations. Colored dots represent trial-averaged individual results, black dots and error bars indicate the grand-average and bootstrapped 95% confidence intervals. Neural tracking of attended speech is stronger under selective attention (C, right plot). Neural tracking of ignored speech is modulated by two-way interactions of spatial cue × semantic cue, and spatial cue × probed ear (D, right plots). Colored dots represent trial-averaged correlation coefficients per participant. Error bars show bootstrapped 95% confidence intervals. The following figure supplements are available for figure 4:

**Figure supplement 1.** Training and testing of envelope reconstruction models.
**Figure supplement 2.** Encoding of attended and ignored speech shown separately per probed-ear condition.
**Figure supplement 3.** Neural tracking strength per reconstruction model and attention condition.
**Figure supplement 4.** Decoding accuracy in selective-attention trials.
**Figure supplement 5.** Decoding accuracy in divided-attention trials.

Figure 4B shows the topographical distribution of regression weights obtained from the linear backward models temporally aligned with the most prominent deflections in the forward-transformed temporal response functions (TRFs) of attended and ignored speech. We averaged models across all N=76 participants, and across attend-left (i.e., ignore-right) and attend-right (i.e., ignore-left) trials (see Fig. 4–supplement 2 for model results split by probed ear).

Both representations show a similar encoding of the attended and ignored sentences in very early stages of processing (< 150ms) as indicated by comparable regression weights over fronto-central channels and corresponding TRF deflections in the time window covering the N1_TRF_ component. By contrast, during ensuing processing stages reflected by the P2_TRF_ component, there are clear differences observable in the strength of neural tracking: For the attended sentence, we observed a pronounced P2_TRF_ component and the absence thereof for the ignored sentence. In addition, in later processing stages (> 300ms), deflections in the TRF for attended and ignored speech follow trajectories of opposite polarity and the associated topographical maps of backward model weights suggest the involvement of distinct brain networks. A complementary analysis examining models results split by probed ear, revealed that this pattern was particularly pronounced in attend-left/ignored-right trials (see Fig. 4- supplements 2A, D).

### Listening cues influence the tracking of attended and ignored speech

To gain a more differentiated understanding of how selective attention and semantic predictability influence the selective cortical representation of speech signals at the single-trial level, we focused on the neural tracking strength of the attended and ignored sentence. More specifically, our linear models predicted the strength of single-trial correlations between the reconstructed and presented envelopes (see Fig. 4–supplement 3 for neural tracking strength per reconstruction model and attention condition).

For selective-attention trials, we observed a degree of neural tracking (attended: mean *r*=.11, range:.003-.22; ignored: mean *r*=.09, range:.007-.23) that was similar in magnitude to that reported in recent neural speech tracking studies using linear backward models (e.g., O’Sullivan et al., 2014; Crosse et al., 2015). However, as shown in Figure 4C and D, the neural tracking of attended and ignored sentences varied across cue-cue combinations. We statistically assessed these cue-driven modulations in two separate linear mixed-effects models (see Tables S5 and S6 for full model details). Note that for sake of consistency, for divided-attention trials, we refer to the reconstruction of the *probed* envelope as the *attended* envelope, and to the reconstruction of the *unprobed* envelope as the *ignored* envelope despite the absence of corresponding instructions.

Giving credence to the suggested role of selective neural speech tracking as a neural filter strategy, we found stronger neural tracking of the attended envelope following an informative spatial-attention cue as compared to an uninformative one (*β*=.08, *p*=.002). The semantic predictability of the sentence-final words, however, did not modulate the neural tracking of the attended envelope (likelihood ratio test *X*^2^_1_=.69, *p*=.41).

Notably, the analysis of ignored-speech tracking revealed a qualitatively different pattern of results: Here, neural tracking strength was modulated by the interactive effect of the spatial-attention cue together with either the semantic cue or the probed ear (spatial cue × semantic cue: *β*=−.12, *p*=.02; spatial cue × probed ear: *β*=−.08, *p*=.006). Following an informative spatial-attention cue, we found increased neural tracking of the ignored sentence when less prior information about the upcoming input was available, that is, when it was preceded by an uninformative semantic cue (simple effect semantic cue, informative spatial cue: *β*=−.08, *p*=.01; uninformative spatial cue: *β*=.03, *p*=.36). At the same time, when participants had to pay attention to both sides, task-irrelevant sentences played to the left ear were tracked more strongly than task-irrelevant sentences played to the right ear (simple effect probed ear, selective: *β*=.04, *p*=.13; divided: *β*=.12, *p* <.001).

### Decoding accuracy

Given that our challenging listening task presented two concurrent sentences that were (i) of short duration, (ii) spoken by the same female talker, (iii) highly similar with respect to their onset envelopes, and (iv) presented against speech-shaped noise, we wished to further evaluate the potency of the employed envelope reconstruction approach under such difficult conditions. To this end, we examined the decoding accuracy across participants in selective-attention trials (see also Fig. 4–supplements 1, 4 and 5). Evaluated at the single-trial level, attention was correctly decoded if the envelope reconstructed by the attended reconstruction model was more similar to the attended envelope than to the ignored envelope (r_attended_ > r_ignored_) and vice versa for the ignored reconstruction model.

Unsurprisingly, the achieved decoding accuracy was lower than that reported in speech tracking studies using much longer segments of continuous speech but still reached a mean decoding accuracy of 55% (range: 0.43–0.65) for the reconstruction model of the attended envelope and 48% (range: 0.36–0.72) for the reconstruction model of the ignored envelope (see Etard et al., 2018 for decoding accuracy as a function of stimulus duration). Importantly, when investigated at the single-subject level, the attended reconstruction model yielded decoding accuracies that exceeded the empirical chance level of 58% in 20 out of 76 participants. The ignored reconstruction model yielded significantly above chance performance in 10 out 76 participants. In addition to the aforementioned experimental conditions that created a particularly difficult scenario for stimulus reconstruction, it is important to bear in mind that decoding accuracy is not only influenced by specifics of the linear model itself but also by how closely participants followed the spatial-cue instructions. For these reasons, our analyses capitalized on the single-trial neural tracking strength as a more direct and fine-grained measure of how strongly the attended and ignored sentence were cortically represented in a particular trial.

### Additional influences on behavioral performance and neural measures

Our modelling approach accounted for a number of additional factors known to challenge speech comprehension, and to modulate neural signature of attention (see Fig. 1). We observed that participants’ performance varied in line with the well-attested *right-ear advantage* (REA, also referred to as *left-ear disadvantage*) in the processing of linguistic materials (Kimura, 1961; Broadbent and Gregory, 1964). More specifically, participants responded both faster and more accurately (response speed: *β*=.06, *p*<.001; accuracy: *OR*=1.38, *p*<.001) when they were probed on the last word presented to the right compared to the left ear. While we observed differences in the strength of ignored-speech tracking across probed-left and probed-right trials under the more challenging divided-attention condition (see section above), the degree of alpha power lateralization did not change with the probed ear (*β*=.03, *p*=.66).

With respect to age and hearing acuity, we observed that decreases in hearing acuity (PTA) led to less accurate response (*OR*=.75, *p*=.007), whereas increased age only led to overall slower (*β*=−.12, *p*=.003) but not less accurate responses (*OR*=.89, *p*=.29). In addition, and in accordance with previous reports (Presacco et al., 2016b), we observed a significant increase in neural tracking of the attended sentence for older participants (*β*=.06, *p*=.02). Again, the strength of alpha power lateralization did not co-vary with participants’ age (*β* =−.01, *p*=.39), confirming results of a previous dichotic listening EEG study on a subsample of the present cohort (Tune et al., 2018).

Lastly, the neural tracking of the attended envelope was stronger when its presentation began prior to that of the ignored envelope (*β*=.33, *p*<.001), which may indicate that the earlier onset of a sentence increased its bottom-up salience and attentional capture.

### Independent single-trial modulation of alpha power lateralization and neural tracking

Our third major goal was to investigate whether the modulation of alpha power and neural tracking strength reflect two dependent neural strategies of attention at all. To this end, using two single-trial linear mixed-effects models, we tested whether the neural tracking of the attended and ignored sentence could be predicted from alpha power dynamics (here, the alpha power at electrodes ipsi- and contralateral to the focus of attention/probed ear, that is, the relative signal modulations that constitute the single-trial alpha lateralization index). We reasoned that it is the up-regulation of contralateral alpha power in the service of selective inhibition, and the down-regulation of ipsilateral alpha power in the service of controlled amplification that in turn might facilitate the differential tracking of attended and ignored speech (cf. Fig. 5A).

**Figure 5.**
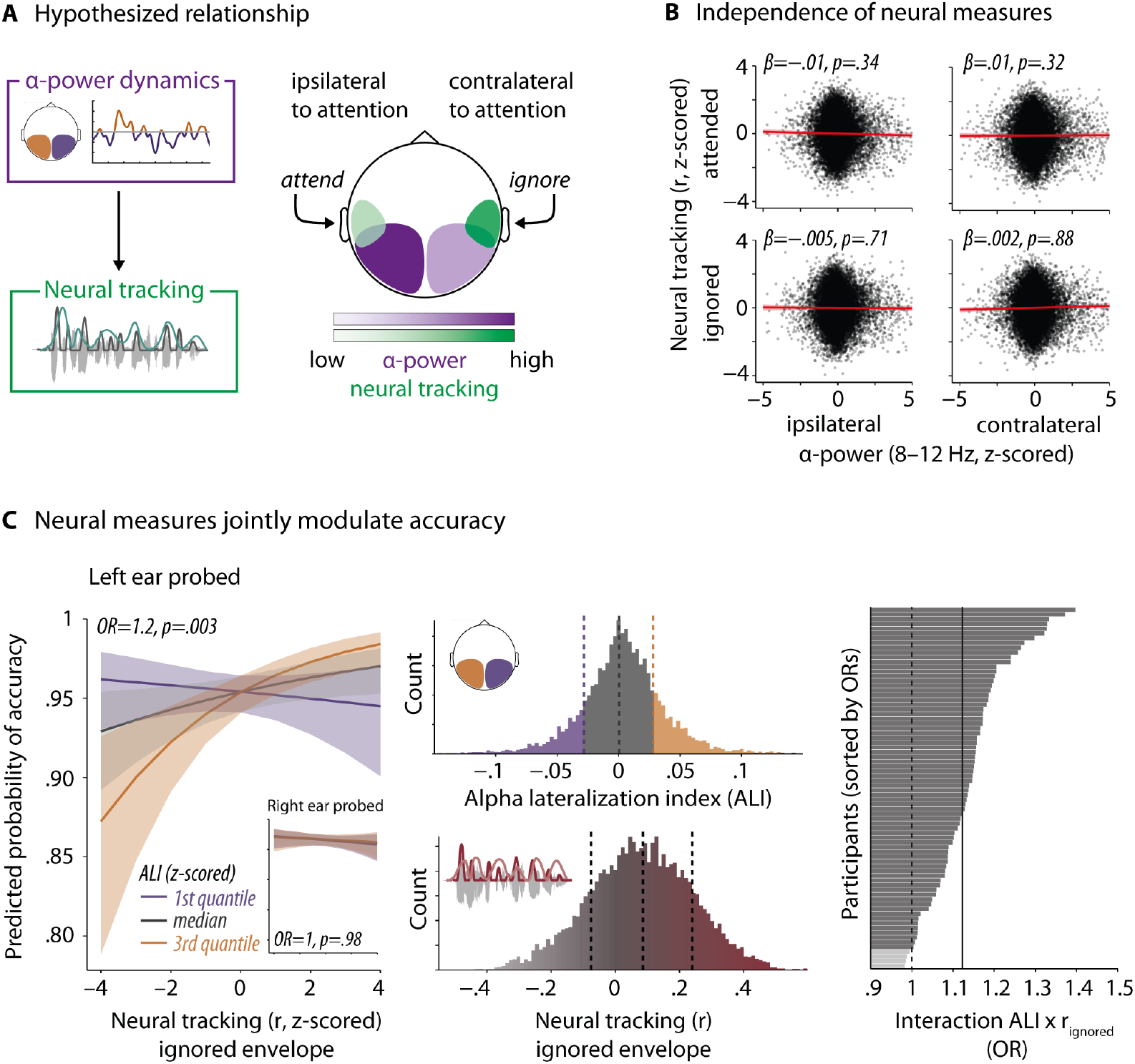
Relationship of alpha power dynamics and neural tracking and their synergistic influence on behavior. (A) Hypothesized inverse relationship of changes in alpha power and neural tracking in the hemisphere contra- and ipsilateral to the focus of attention. Changes in alpha power are assumed to drive changes in neural tracking strength. Schematic representation of expected co-variation in the two neural measures during an attend-left trial. (B) Independence of changes in neural tracking and alpha power during sentence processing as revealed by two separate linear mixed-effects models predicting the neural tracking strength of attended and ignored speech, respectively. Error band denotes 95% confidence interval. Black dots represent single-trial observed values. (C, left panel) Visualization of the interactive effect of alpha power lateralization (ALI) and neural tracking of the ignored envelope as predicted from the generalized linear model for probed-left trials. Inset shows absence of the effect in probed-right trials. Neural measures are z-scored across all N=76 participants. The relationship of neural tracking and accuracy is shown for three quantiles of the ALI. Error bars and bands represent 95% confidence intervals. (C, middle panel) Distribution of ALI (top) and neural tracking raw values (bottom) prior to z-scoring. Vertical dashed lines index the median, first and third quantiles. (C, right panel) Consistency of the two-way interaction of ALI and neural tracking across all 76 participants. Effects for individual participants are estimated via the inclusion of corresponding random slopes in a model of probed-left trials, only. Solid black line represents the fixed-effects level interaction effect; black dotted line indicates an odds ratio (OR) of 1. Dark grey bars are individual subjects with an OR above 1; light grey bars show subjects with an OR of below 1. The following figure supplements are available for figure 5:

**Figure supplement 1.** Modulation of response speed by neural tracking strength (r_attended_) per attention condition.

However, neither the tracking of the attended envelope (main effect alpha ipsi: *β*=−.01, *p*=.36; main effect alpha contra: *β*=.01, *p*=.32; interaction ipsi × contra: *β*=−.001, *p*=.54) nor the tracking of the ignored envelope could be predicted by changes in ipsi- or contralateral alpha power or by their interaction (main effect alpha ipsi: *β*=−.005, *p*=.71; main effect alpha contra: *β*=.002, *p*=.88; interaction ipsi × contra: *β*=−.001, *p*=.64; see Tables S6 and S7 for full model details).

This notable absence of an alpha lateralization-speech tracking relationship was consistent across the spatial-attention and probed-ear conditions (i.e., inclusion of the respective terms did not improve model fit; all *ps* > 0.2). Importantly, the observed independence of these two neural measures also did not hinge on whether we modeled the change in alpha power as two separate regressors or included the strength of the alpha lower lateralization expressed by the ALI instead (see Table S9 for details).

### Neural strategies of attention jointly support listening success under challenging conditions

Having established the surprising functional independence of alpha power lateralization and speech tracking, the final and most important piece of our investigation becomes in fact statistically more tractable: If alpha power dynamics in parieto-occipital cortex, and neural speech tracking in auditory cortices essentially act as two independent neural filter strategies, we can proceed to probe their relative functional relevance for behavior in a factorial-design fashion. That is, we were interested in how far changes in listening success related to the independent (i.e., as main effects) or joint influence (i.e., as an interaction) of neural measures. We answered these question using the same linear mixed-effects models as in testing our first research question (Q1 in Fig.1), as well as the influence of additional factors (see Tables S1 and S2 for details).

First, neural tracking of the attended envelope had opposing effects on listeners’ response speed, depending on the attention condition (interaction neural tracking × spatial cue: *β*=−.04, *p*=.004J: In the behaviorally more difficult divided-attention condition, increased tracking of the to-be-attended sentence led to faster responses (*β*=.02, *p*=.01J. In cued selective-attention trials, we found only a weaker trend, in the opposite direction (*β*=−.02, *p*=.1; see Fig. 5- supplement 1).

Secondly, for response accuracy as our most important indicator of listening success, we also found a beneficial effect of stronger neural tracking that occurred under the behaviorally more challenging condition: Specifically, we observed a joint influence of the degree of alpha power lateralization and ignored-speech tracking when the less favorable left ear was probed (interaction neural tracking ignored × ALI × probed ear: *OR*=.86, *p*=.03; simple effects neural tracking ignored – left ear: *OR=1*.17, *p*=.003; right ear: *OR=1, p*=.98).

Probed-left trials substantially increased the probability of giving a correct answer for neural states of co-occurring strong alpha power lateralization and strong ignored-speech tracking (Fig. 5C). This particular brain-behavior relationship was highly consistent across the large sample of participants, and we observed a corresponding effect in 72 out of 76 participants.

Moreover, this joint influence of alpha power lateralization (ALI) and neural tracking was specific to their joint occurrence in the time window of sentence presentation. We did not observe any comparable relationship of ALI and neural tracking on behavior when we substituted the degree of alpha power lateralization during sentence presentation for the degree of lateralization during the spatial-attention cue (see Tables S10 and S11 for details).

## Discussion

In the present study, we utilized the power of a large, representative sample of middle-aged and older listeners to explicitly address the question of how two different neural filter strategies, typically studied in isolation, jointly shape listening success. In our dichotic listening task, we recorded the electroencephalogram and observed systematic changes in alpha power lateralization and the neural tracking of attended speech in response to an informative spatial-attention cue. These results not only provide unprecedented large-sample support for their suggested roles as neural signatures of selective attention. We also were able to address two important but still insufficiently answered questions: How do the two neural filter strategies relate to one another, and how do they impact speech comprehension success in a demanding real-life listening situation?

First, when related at the single-trial, single-subject level, we found the modulation of the two neural measures to be statistically independent, underlining their functional segregation and speaking to two distinct neurobiological implementations.

Secondly, the presence of an informative spatial-attention cue not only boosted both neural measures of interest but also consistently boosted behavioral performance. However, changes at the neural and at the behavioral levels were not generally related. Instead, we observed a joint influence of alpha power lateralization and ignored-speech tracking on the behavioral outcome only for trials in which the slightly speech-disadvantaged left ear was probed.

### Alpha power and neural speech tracking implement independent neural filter strategies

Attention-related changes in alpha power dynamics and in the neural tracking of speech signals arise from distinct cortical areas; involve neurophysiological signals operating at different frequency regimes; and are typically investigated in separate lines of research. Nevertheless, there is preliminary evidence suggesting that these two neural filter strategies may exhibit a systematic relationship (Kerlin et al., 2010; Lakatos et al., 2016; Wöstmann and Obleser, 2016; Henry et al., 2017; Tune et al., 2018). Therefore, in the present study we probed the hypothesis that a domain-general, top-down controlled attentional filter implemented via regulation of alpha power in parieto-occipital cortices would in turn facilitate the selective sensory analysis of speech signals in auditory cortex (Schroeder and Lakatos, 2009; Schroeder et al., 2010).

However, our analyses of trial-by-trial variation in alpha power and neural tracking of attended and ignored speech revealed that the relative up- or down-regulation of alpha power varied independently of the degree to which the attended or the ignored sentence were neurally tracked in auditory cortex. Therefore, the present results speak against a consistent, linear relationship of neural filter strategies. We see instead the coexistence of two complementary but independent neural solutions to the implementation of auditory selective attention for the purpose of speech comprehension.

The proposed independence of neural filters in the current task is supported by additional pieces of evidence: On the one hand, the two neural measures were differentially affected by additional factors modelled in our analyses: The strength of alpha power lateralization was only influenced by the spatial-attention cue. By contrast, and in line with previous reports, the degree of neural tracking of attended and ignored speech varied in line with both the spatial-attention and semantic cue, participants’ age, as well as the probed ear (Sohoglu et al., 2012; Peelle et al., 2013; Presacco et al., 2016a; 2016b). These results corroborate the assumption of separate cortical circuits of distinct bottom-up and top-down communication that give rise to the neural filter strategies under investigation here.

Further support stems from a recent fMRI study from our laboratory that used the same dichotic listening task and tested a subsample of the present participant cohort (Alavash et al., 2018). In a graph-theoretic analysis of cortex-wide networks, the reconfiguration of resting state networks in adaptation to the listening task did not increase integration of larger brain networks. Instead it prioritized local processing by means of higher segregation. Crucially, switching from rest to task triggered the emergence of an auditory control sub-network that enabled a fine-tuned cross-talk between auditory, temporal, and frontal control areas. That is, we observed a reconfigured composition of brain networks that involved brain areas generally associated with the neural tracking of speech but not those typically engaged in the controlled inhibition of irrelevant information via top-down regulation of alpha power (Banerjee et al., 2011; Nourski and Brugge, 2011; Giraud and Poeppel, 2012; Sadaghiani and Kleinschmidt, 2016).

What can be concluded about the functional interplay of the two neural filter strategies when put in context of previous electrophysiological studies reporting diverging results (Kerlin et al., 2010; Lakatos et al., 2016; Henry et al., 2017)? Taken together, the results do provide strong evidence in favor of a generally existent functional tradeoff between attentional filter mechanisms. However, we suggest that the precise nature of this interplay of attentional filter mechanisms hinges on a number of factors such as the particular task demands, and potentially on the level of temporal and/or spatial resolution at which the two neural signatures are examined (Lakatos et al., 2016).

### Synergistic interaction of neural filter strategies supports speech comprehension under challenging conditions

With the present study, we explicitly addressed the often overlooked question of how trial-by-trial changes at the neural level would impact behavior, here single-trial speech comprehension success (van Ede et al., 2012; Ding and Simon, 2014; Krakauer et al., 2017). Notably, despite an unprecedentedly large sample of almost eighty listeners, adequate number of within-subject trials, and a sophisticated linear-model approach, we did not find evidence of a direct (i.e. independent of the listening cues or related factors such as the probed ear) influence of the relative strength of alpha power lateralization or of the neural tracking of the attended or ignored sentence.

Instead, for trials in which the left ear was probed, we observed a significant modulation of accuracy that depended on the relative levels of both alpha power lateralization and the neural tracking of the ignored sentence. Highly consistent across individuals, in neural states of strong alpha lateralization a sizable performance difference arises from the degree of neurally tracking the ignored speech signal: There was a predicted 20-% increased chance of accurately responding in a trial, as ignored-speech tracking moved from weakest to strongest.

In light of the well-established right ear advantage (Kimura, 1961; Broadbent and Gregory, 1964), observing such an intricate interaction pattern should come as no surprise to psychoacousticians, psychologists, and neuroscientists alike. Nevertheless, the dependence on probed ear is an exploratory finding and begs replication in the second half of our large-scale, ongoing cohort acquisition (intended total N is 160).

Despite the exploratory nature of this last finding, the elaborate circumstances under which the significant influence of neural changes on behavior became manifest, highlight the complexity of often implicitly assumed brain-behavior relationships and their dependence on a multitude of contextual factors. Moreover, even in cases where a consistent relationship of attention-related neural fluctuations and a behavioral outcome can be established, the neural contribution to behavioral change may be surprisingly low (as shown for alpha power modulation before; van Ede et al., 2012).

Indeed, in previous studies on alpha power lateralization, in particular those that focused on middle-aged and older adults, evidence for the behavioral importance of this neural signature has been mixed (Sander et al., 2012; Hong et al., 2015; Mok et al., 2016; Leenders et al., 2018). The present results are generally in line with one of our own recent studies carried out on a subsample of the present cohort. Using a fine-grained analysis of concurrent neural and behavioral changes, we found only a weak link between the strength of alpha lateralization and accuracy in a similar dichotic listening task (Tune et al., 2018).

Given the research field’s present focus on highly naturalistic designs (Hamilton and Huth, 2018), such as the concurrent presentation of two narratives, evidence on the functional importance of selective neural speech tracking is generally sparse (Power et al., 2012; Zion Golumbic et al., 2013; O’Sullivan et al., 2014; Fiedler et al., 2017; 2019). A notable exception is the study by Mesgarani and Chang (2012) who observed an enhanced auditory cortical representation of the attended versus ignored talker in correct compared to error trials. Further supporting the idea that the behavioral importance of neural filter strategies may depend on the level of task difficulty, overall performance in this task was notably lower (~70%) than in the present study.

Consequently, a testable hypothesis arises: The more attentionally demanding a listening situation is, the more will listening success depend on the fidelity of neural filter strategies and their potential interaction. The present data hold initial evidence for this assumption: The overall good behavioral performance (around 90%) attests that our task had a moderate level of difficulty. The joint influence of the lateralized alpha power and ignored-speech speech tracking on accuracy occurred only under the relatively more demanding condition, that is, when the behaviorally disadvantaged left ear was probed.

### Neural tracking of irrelevant speech is not irrelevant to listening success

A crucial finding of the present study lies in the behavioral importance of the degree to which the ignored sentence was neurally tracked. This finding adds to a growing body of evidence suggesting that the successful deployment of auditory selective attention depends on the neural fate of both to-be-attended and to-be-ignored speech signals (e.g. Melara et al., 2002; Chait et al., 2010; for review see Shinn-Cunningham, 2008, Shinn-Cunningham and Best, 2008; Kaya and Elhilali, 2017).

Our results are generally compatible with the findings of a recent, source-localized EEG study (Fiedler et al., 2019) providing evidence for the active suppression of to-be-ignored speech signals via an enhanced cortical representation of the ignored talker. Importantly, this amplification in the neural representation of to-be-ignored speech in areas beyond auditory cortex was only found in situations where the distracting talker was louder and thus perceptually more dominant.

In absence of any systematic manipulation of the signal-to-noise ratio (SNR) of attended and ignored speech, the trial-by-trial differences in the probed ear employed here may be likened to a shift in the subjectively perceived SNR. That is, the observed right ear benefit for both accuracy and response speed may in part be driven by the greater difficulty of tuning out the to-be-ignored sentence when it was played to the right compared to the left ear.

Our data provide preliminary support for this assumption of a differential neural representation of ignored speech presented to the right versus the left ear (see Fig. 4–supplements 2A, D): In attend-left/ignore-right trials, we observed more selective cortical representations of attended and ignored speech, crucially supported by a late distinct response to the ignored speech signal. While additional research should systematically test the hypothesis put forward here, our results suggest that under relatively challenging listening situation (e.g., when the behaviorally disadvantages left ear is probed), listening success increasingly depends on the selective neural encoding of both attended and ignored speech.

## Conclusion

In a large, representative sample of adult listeners, we have provided evidence that single-trial listening success in a challenging, dual-talker acoustic environment will not be meaningfully modelled when focusing on a single, one-to-one neural substrate alone: An attentional cue increases the engagement of two distinct neural filter strategies and boosted listening success. However, there was no direct link between behavioral and neural changes. Instead, the observed joint influence of alpha power lateralization and ignored-speech tracking in trials where the slightly speech-disadvantaged left ear was probed highlights the intricate nature of this brain-behavior relationship and should temper over-simplified accounts of the predictive power of neural filter strategies for behavior. It also emphasizes a third major finding of this study, potentially disruptive to ongoing research programs in decoding attentional states from the listening brain: There is behavioral relevance to the degree of neurally representing ignored speech in the cortical response.

## Materials and Methods

### Participants and procedure

Seventy-six right-handed German native speakers (median age 56 years; range 39–70 years; 28 males; see Fig. 1–supplement 1 for age distribution) were included in the sample. All participants are part of a large-scale study on the neural and cognitive mechanisms supporting adaptive listening behavior in healthy middle-aged and older adults (“The listening challenge: How ageing brains adapt (AUDADAPT)”; https://cordis.europa.eu/project/rcn/197855_en.html). Handedness was assessed using a translated version of the Edinburgh Handedness Inventory (Oldfield, 1971). All participants had normal or corrected-to-normal vision, did not report any neurological, psychiatric, or other disorders and were screened for mild cognitive impairment using the German version of the 6-Item Cognitive Impairment Test (6CIT; Jefferies and Gale, 2013).

As part of our large-scale study on adaptive listening behavior in healthy aging adults, participants also underwent a session consisting of a general screening procedure, detailed audiometric measurements, and a battery of cognitive tests and personality profiling (see Tune et al., 2018 for details). This session always preceded the EEG recording session. Only participants with normal hearing or age-adequate mild-to-moderate hearing loss were included (see Fig. 1–supplement 2 for individual audiograms). Participants gave written informed consent and received financial compensation (8€ per hour). Procedures were approved by the ethics committee of the University of Lübeck and were in accordance with the Declaration of Helsinki.

### Dichotic listening task

In a novel linguistic variant of a classic Posner paradigm (Posner, 1980), participants listened to two competing, dichotically presented sentences. They were probed on the sentence-final word in one of the two sentences. Critically, two visual cues preceded auditory presentation. First, a spatial-attention cue either indicated the to-be-probed ear, thus invoking selective attention, or did not provide any information about the to-be-probed ear, thus invoking divided attention. Second, a semantic cue specified a general or a specific semantic category for the final word of both sentences, thus allowing to utilize a semantic prediction. Cue levels were fully crossed in a 2×2 design and presentation of cue combinations varied on a trial-by-trial level (Fig. 2A). The trial structure is exemplified in Figure 2B. Details on stimulus construction and recording can be found in the Supplemental Information.

Each trial started with the presentation of a fixation cross in the middle of the screen (jittered duration: mean 1.5 s, range 0.5–3.5 s). Next, a blank screen was shown for 500 ms followed by the presentation of the spatial cue in the form of a circle segmented equally into two lateral halves. In selective-attention trials, one half was black, indicating the to-be-attended side, while the other half was white, indicating the to-be-ignored side. In divided-attention trials, both halves appeared in grey. After a blank screen of 500 ms duration, the semantic cue was presented in the form of a single word that specified the semantic category of both sentence-final words. The semantic category could either be given at a general (natural vs. man-made) or specific level (e.g. instruments, fruits, furniture) and thus provided different degrees of semantic predictability. Each cue was presented for 1000 ms.

After a 500 ms blank-screen period, the two sentences were presented dichotically along with a fixation cross displayed in the middle of the screen. Finally, after a jittered retention period, a visual response array appeared on the left or right side of the screen, presenting four word choices. The location of the response array indicated which ear (left or right) was probed. Participants were instructed to select the final word presented on the to-be-attended side using the touch screen. Among the four alternative were the two actually presented nouns as well as two distractor nouns from the same cued semantic category.

Stimulus presentation was controlled by PsychoPy (Peirce, 2007). The visual scene was displayed using a 24” touch screen (ViewSonic TD2420) positioned within an arm’s length. Auditory stimulation was delivered using inear headphones (EARTONE 3A) at sampling rate of 44.1 kHz. Following instructions, participants performed a few practice trials to familiarize themselves with the listening task. To account for differences in hearing acuity within our group of participants, individual hearing thresholds for a 500-ms fragment of the dichotic stimuli were measured using the method of limits. All stimuli were presented 50 dB above the individual sensation level. During the experiment, each participant completed 60 trials per cue-cue condition, resulting in 240 trials in total. The cue conditions were equally distributed across six blocks of 40 trials each (~ 10 min) and were presented in random order. Participants took short breaks between blocks.

### EEG data acquisition and preprocessing

Participants were seated comfortably in a dimly-lit, sound-attenuated recording booth where we recorded their EEG from 64 active electrodes mounted to an elastic cap (Ag/AgCl; ActiCap / ActiChamp, Brain Products, Gilching, Germany). Electrode impedances were kept below 10 kΩ. The signal was digitized at a sampling rate of 1000 Hz and referenced on-line to the left mastoid electrode (TP9, ground: AFz).

For subsequent off-line EEG data analyses, we used the EEGlab (version 14_1_1b; Delorme and Makeig, 2004) and Fieldtrip toolboxes (version 2016-06-13; Oostenveld et al., 2011), together with customized Matlab scripts. Independent component analysis (ICA) using EEGlab’s default runica algorithm was used to remove all non-brain signal components including eye blinks and lateral eye movements, muscle activity, heartbeats and single-channel noise. Prior to ICA, EEG data were re-referenced to the average of all EEG channels (average reference). Following ICA, trials during which the amplitude of any individual EEG channel exceeded a range of 200 microvolts were removed.

### Behavioral data analysis

We evaluated participants’ behavioral performance in the listening task with respect to accuracy and response speed. For the binary measure of accuracy, we excluded trials in which participants failed to answer within the given 4-s response window (‘timeouts’). Spatial stream confusions, that is trials in which the sentence-final word of the to-be-ignored speech stream were selected, and random errors were jointly classified as incorrect answers. The analysis of response speed, defined as the inverse of reaction time, was based on correct trials only. Single-trial behavioral measures were subjected to (generalized) linear mixed-effects analysis (see Statistical analysis).

### EEG data analysis

#### Time-frequency analysis

To obtain time-frequency representations of single-trial oscillatory power, the preprocessed continuous EEG data were high-pass-filtered at 0.3 Hz (finite impulse response (FIR) filter, zero-phase lag, order 5574, Hann window) and low-pass-filtered at 180 Hz (FIR filter, zero-phase lag, order 100, Hamming window). The EEG was cut into epochs of -2 to 8 s relative to the onset of the spatial-attention cue to capture cue presentation as well as the entire auditory stimulation interval. Individual epochs were convolved with frequency-adaptive Hann-tapers of 7 cycles width (2-30 Hz, step size 0.5 Hz). Oscillatory power was estimated by squaring the magnitude of the complex-valued Fourier coefficients at each single frequency. Since oscillatory power or amplitude values typically follow a highly skewed, non-normal distribution, we applied a nonlinear transformation of the Box-Cox family (power_*trans*_ = (power^p^ -1)/p with *p*=0.2) to minimize skewness and to satisfy the assumption of normality for parametric statistical tests involving oscillatory power values (Smulders et al., 2018).

To capture whole-trial (0-7 s) dynamics in oscillatory power across all trials, we first subtracted power averaged across trials in this interval from individual trials (absolute baseline). Grand average representations of oscillatory power were obtained by averaging the result across all trials and participants.

#### Attentional modulation of alpha power

To quantify attention-related changes in 8-12 Hz alpha power, we calculated two attentional modulation indices separately for selective- and divided-attention trials: the topographical alpha lateralization index [ALI_topo_ = (α-power_att-left_ – α-power_att-right_) / (α-power_att-left_ + α-power_att-right_)] and the single-trial, temporally resolved alpha lateralization index [ALI = (α-power_ipsi_ – α- power_contra_)/(α-power_ipsi_ + α-power_contra_)] (Haegens et al., 2011; Wöstmann et al., 2016; Tune et al., 2018).

The ALI_topo_, calculated per channel for 8-12 Hz absolute alpha power, provides a spatial representation of attentional effects. Positive ALI_topo_ values reflect higher alpha power levels during attend-left compared to attend-right trials. The reverse relation holds for negative values. For divided attention trials in which attention was not cued to one of the two sides, trials were categorized based on the probed ear.

In estimating the ALI_topo_ we used a robust variant of the index that applies the inverse logit transform [(1 / 1+ exp(-x)] to both inputs to scale them into a common, positive-only [0;1]-bound space prior to index calculation.

To calculate the ALI, which provides time-resolved single-trial estimates of the attentional modulation of alpha power, we first corrected for overall hemispheric power differences independent of attentional modulation. To this end, we normalized single-trial power values by calculating per channel and frequency the whole-trial (0-7 s) power averaged across all trials and subtracting it from single trials (see Fig. 3–supplement 2 for ALI calculated on absolute power values).

We then selected per participant the ten channels with the largest positive ALI_topo_ values over the left posterior hemisphere and the ten channels with the largest negative ALI_topo_ values over the right posterior hemisphere. For each of the two attention conditions (attend-/probed-left versus attend/probed-right), the selected channels were classified as being either ipsi- or contralateral to the focus of attention (in divided-attention trials: probed ear). To obtain a time-resolved measure of attentional modulation of alpha power, we calculated the ALI over the time course of the entire trial using a 250-ms sliding window (Mesgarani and Chang, 2012). Index calculation applied the same inverse logit transformation as used for the ALI_topo_. For statistical analysis of cue-driven neural modulation, we extracted single-subject mean ALI values per experimental condition in the time window of sentence presentation (3.5-6.5 s), and treated them as the dependent measure in linear mixed-effects analysis. In addition, they served as continuous predictors in the statistical analysis of brain-behavior relationships (see below).

#### Neural speech tracking analysis

To assess the neural encoding of speech by low-frequency activity, the continuous preprocessed EEG were down-sampled to f_s_=125 Hz, filtered between f_c_=1 and 8 Hz (FIR filters, zero-phase lag, order: 8f_s_/f_c_ and 2f_s_/f_c_, Hamming window), and re-referenced to the average of both mastoid electrodes (TP9 and TP10). The EEG was cut to yield individual epochs covering the presentation of auditory stimuli, beginning at noise onset until the end of auditory presentation. Individual epochs were z-scored prior to submitting them to regression analysis (see below).

#### Extraction of envelope onsets

From the presented speech signals, we derived a temporal representation of the acoustic onsets in the form of the onset envelope (cf. Fiedler et al., 2019). To this end, using the NSL toolbox (Chi et al., 2005), we first extracted an auditory spectrogram of the auditory stimuli (128 spectrally resolved sub-band envelopes logarithmically spaced between 90-4000 Hz), which were then summed across frequencies to yield a broad-band temporal envelope. Next, the output was down-sampled and low-pass filtered to match the specifics of the EEG. To derive the final onset envelope to be used in linear regression, we first obtained the first derivative of the envelope and set negative values to zero (half-wave rectification) to yield a temporal signal with positive-only values reflecting the acoustic onsets (see Fig. 4A and figure supplements).

#### Estimation of envelope reconstruction models

To investigate how low-frequency (i.e, < 8 Hz) fluctuations in EEG activity relate to the encoding of attended and ignored speech, we used a linear regression approach to describe the mapping from the presented speech signal to the resulting neural response (Lalor and Foxe, 2010; Crosse et al., 2016). More specifically, we trained stimulus reconstruction models (also termed decoding or backward models) to predict the onset envelope of the attended and ignored speech stream from EEG. In this analysis framework, a linear reconstruction model *g* is assumed to represent the linear mapping from the recorded EEG signal, *r*(*t,n*), to the stimulus features, *s*(*t*):

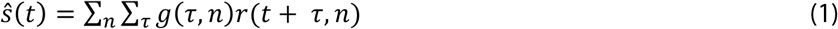

where *s^*(*t*) is the reconstructed onset envelope at time point *t*. We used all 64 EEG channels and time lags τ in the range of -100 ms to 500 ms to compute envelope reconstruction models using ridge regression (Hoerl and Kennard, 1970):

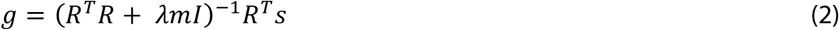

where *R* is a matrix containing the sample-wise time-lagged replication of the neural response matrix *r*, λ is the ridge parameter for regularization, *m* is a scalar representing the mean of the trace of R^T^R (Biesmans et al., 2016), and *I* is the identity matrix. We followed the procedures described in (Fiedler et al., 2017) to estimate the optimal ridge parameter at λ=5.

Compared to linear forward (‘encoding’) models that derive temporal response functions (TRFs) independently for each EEG channel, stimulus reconstruction models represent multivariate impulse response functions that exploit information from all time lags and EEG channels simultaneously. To allow for a neurophysiological interpretation of backward model coefficients, we additionally transformed them into linear forward model coefficients following the inversion procedure described in Haufe et al. (2014). All analyses were performed using the multivariate temporal response function (mTRF) toolbox (version 1.5; Crosse et al., 2016) for Matlab.

Prior to model estimation, we split the data based on the two spatial attention conditions (selective vs. divided), resulting in 120 trials per condition. Envelope reconstruction models were trained on selective-attention single-trial data, only. Two different backward models were estimated for a given trial, an envelope reconstruction model for the-be-attended speech stream (short: attended reconstruction model), and one for the to-be-ignored speech stream (short: ignored reconstruction model). Reconstruction models for attended and ignored speech signals were trained separately for attend-left and attend-right trials which yielded 120 single-trial decoders (60 attended, 60 ignored) per attentional setting. For illustrative purposes and to evaluate decoding accuracy, we averaged the reconstruction models for attended and ignored envelopes and their forward transformations across left and right trials for all participants (Fig. 4B).

#### Evaluation of envelope reconstruction accuracy

To quantify how strongly the attended and ignored sentences were tracked by slow cortical dynamics, at the single-subject level, we reconstructed the attended and ignored envelope of a given trial using a leave-one-out cross-validation procedure. Following this approach, the envelopes of each trial were reconstructed using the averaged reconstruction models trained on all but the tested trial. For a given trial, we only used the trained models that corresponded to the respective cue condition (i.e., for an attend-left/ignore-right trial we only used the reconstruction models trained on the respective trials). The reconstructed onset envelope obtained from each model was then compared to the two onset envelopes of the actually presented speech signals. The resulting Pearson-correlation coefficients, r_attended_ and r_ignored_, reflect the single-trial neural tracking strength or reconstruction accuracy (cf. O’Sullivan et al., 2014; see also Fig. 4–supplement 1).

We proceeded in a similar fashion for divided-attention trials. Since these trials could not be categorized based on the to-be-attended and -ignored side, we split them based on the ear that was probed at the end the trial. Given that even in the absence of a valid attention cue, participants might still (randomly) focus their attention to one of the two streams, we wanted to quantify how strongly the probed and unprobed envelopes were tracked neurally. To this end, we used the reconstruction models trained on selective-attention trials to reconstruct the onset envelopes of divided-attention trials. Sentences presented in probed-left/unprobed-right trials were reconstructed using the attend-left/ignore-right reconstruction models while probed-right/unprobed-left trials used the reconstruction models trained on attend-right/ignore-left trials.

One of the goals of the present study was to investigate the degree to which the neural tracking of the attended and ignored envelopes would be influenced by spatial attention and semantic predictability, and how these changes would relate to behavior. We thus used the neural tracking strength of the attended envelope (when reconstructed with the attended reconstruction model) and that of the ignored envelope (when reconstructed with the ignored reconstruction model) as (in-)dependent variables in our linear mixed-effects analyses (see below).

#### Decoding accuracy

To evaluate how accurately we could decode an individual’s focus of attention, we separately evaluated the performance of the reconstruction models for the attended or the ignored envelope. When the reconstruction was performed with the attended reconstruction models, the focus of attention was correctly decoded if the correlation with the attended envelope was greater than that with the ignored envelope. Conversely, for reconstructions performed with the ignore reconstruction models, a correct identification of attention required the correlation with the ignored envelope to be greater than that with the attended envelope (see Fig. 4–supplement 1). To quantify whether the single-subject decoding accuracy was significantly above chance, we used a binomial test with α=0.05.

### Statistical analysis

We used (generalized) linear mixed-effect models to investigate the influence of the experimental cue conditions (spatial cue: divided vs. selective; semantic cue: general vs. specific) on behavior, see Q1 in Fig. 1) as well as on our neural measures of interest (Q2). Finally, we investigated the relationship of neural measures (Q3) and their (joint) influence of the different cue conditions and neural measures on behavior (Q4). Using linear mixed-effects models allowed us to model and control for the impact of various additional covariates known to influence behavior and/or neural measures. These included the probed ear (left/right), whether the later-probed sentence had the earlier onset (yes/no), as well as participants’ age and hearing acuity (pure-tone average across both ears).

Model estimation and selection followed an iterative model fitting procedure that started with an intercept-only null model (Tune et al., 2018). Fixed effects terms were added in a stepwise procedure in the order of their conceptual importance for the tested hypotheses, beginning with main effects followed by higher-order interactions, and the change of model fit (fitted using maximum-likelihood estimation) was assessed using likelihood ratio tests. The effects included in the brain-behavior model were guided by effects found in the cue-driven neural modulation models. Deviation coding was used for categorical predictors. All continuous variables were z-scored.

For the dependent measure of accuracy, we used generalized linear mixed-effects model (binomial distribution, logit link function). For response speed, we used general linear mixed-effects model (gaussian distribution, identity link function). *P*-values for individual model terms in the linear models are based on *t*-values and use the Satterthwaite approximation for degrees of freedom (Luke, 2017). *P*-values for generalized linear models are based on z-values and asymptotic Wald tests. Post hoc linear contrasts for categorical regressors with more than two levels were carried out on predicted marginal means (‘least-square means’) and standard errors estimated from the model (Lenth, 2016). In lieu of a standardized measure of effect size for mixed-effects models, we report odds ratios (OR) for generalized linear models and standardized regression coefficients (β) for linear models.

All analyses were performed in R (R Core Team, 2018) using the packages lme4 (Bates et al., 2015), emmeans (Lenth, 2016) and sjPlot (Lüdecke, 2018).

### Data availability

The complete dataset associated with this work including raw data, EEG data analysis results, as well as corresponding code will be publicly available under https://osf.io/28r57/.

## Acknowledgement

Felix Deilmann, Elisabeth Ni, Franziska Scharata, Philipp Seidel, and Annika Struck helped acquire and manage the data. Research was supported by the European Research Council (ERC Consolidator grant AUDADAPT, no. 646696) to JO.

## References

Ahveninen J, Huang S, Belliveau JW, Chang W-T, Hämäläinen M (2013) Dynamic oscillatory processes governing cued orienting and allocation of auditory attention. J Cogn Neurosci 25:1926–1943.

Alavash M, Tune S, Obleser J (2018) Modular reconfiguration of an auditory-control brain network supports adaptive listening behavior. Proc Natl Acad Sci USA 48:201815321–10.

Banerjee S, Snyder AC, Molholm S, Foxe JJ (2011) Oscillatory Alpha-Band Mechanisms and the Deployment of Spatial Attention to Anticipated Auditory and Visual Target Locations: Supramodal or Sensory-Specific Control Mechanisms? J Neurosci 31:9923–9932.

Broadbent DE, Gregory M (1964) Accuracy of recognition for speech presented to the right and left ears. Quarterly Journal of Experimental Psychology 16:359–360.

Chait M, de Cheveigné A, Poeppel D, Simon JZ (2010) Neural dynamics of attending and ignoring in human auditory cortex. Neuropsychologia 48:3262–3271.

Cherry EC (1953) Some experiments on the recognition of speech, with one and with two ears. J Acoust Soc Am 25:975–979.

Chi T, Ru P, Shamma SA (2005) Multiresolution spectrotemporal analysis of complex sounds. J Acoust Soc Am 118:887–906.

Crosse MJ, Butler JS, Lalor EC (2015) Congruent Visual Speech Enhances Cortical Entrainment to Continuous Auditory Speech in Noise-Free Conditions. J Neurosci 35:14195–14204.

Crosse MJ, Di Liberto GM, Bednar A, Lalor EC (2016) The Multivariate Temporal Response Function (mTRF) Toolbox: A MATLAB Toolbox for Relating Neural Signals to Continuous Stimuli. Front Hum Neurosci 10:604.

Delorme A, Makeig S (2004) EEGLAB: an open source toolbox for analysis of single-trial EEG dynamics including independent component analysis. Journal of Neuroscience Methods 134:9–21.

Ding N, Simon JZ (2012) Emergence of neural encoding of auditory objects while listening to competing speakers. Proc Natl Acad Sci USA 109:11854–11859.

Ding N, Simon JZ (2014) Cortical entrainment to continuous speech: functional roles and interpretations. Front Hum Neurosci 8:13367.

Bates D, Mächler M, Bolker B, Walker S (2015) Fitting Linear Mixed-Effects Models Using lme4. J Stat Soft 67:1–48.

Etard O, Kegler M, Braiman C, Forte AE, Reichenbach T (2018) Real-time decoding of selective attention from the human auditory brainstem response to continuous speech. bioRxiv doi: 10.1101/259853

Fiedler L, Wöstmann M, Graversen C, Brandmeyer A, Lunner T, Obleser J (2017) Single-channel in-ear-EEG detects the focus of auditory attention to concurrent tone streams and mixed speech. Journal of Neural Engineering 14: 036020.

Fiedler L, Wöstmann M, Herbst SK, Obleser J (2019) Late cortical tracking of ignored speech facilitates neural selectivity in acoustically challenging conditions. NeuroImage 186:33–42.

Foxe JJ, Snyder AC (2011) The role of alpha-band brain oscillations as a sensory suppression mechanism during selective attention. Front Psychology 2:154.

Giraud AL, Poeppel D (2012) Cortical oscillations and speech processing: emerging computational principles and operations. Nat Neurosci 15:511–517.

Haegens S, Händel BF, Jensen O (2011) Top-down controlled alpha band activity in somatosensory areas determines behavioral performance in a discrimination task. J Neurosci 31:5197–5204.

Hamilton LS, Huth AG (2018) The revolution will not be controlled: neural stimuli in speech neuroscience. Lang Cogn Neurosci 27:1–10.

Haufe S, Meinecke F, Görgen K, Dähne S, Haynes J-D, Blankertz B, Bieβmann F (2014) On the interpretation of weight vectors of linear models in multivariate neuroimaging. NeuroImage 87:96–110.

Händel BF, Haarmeier T, Jensen O (2011) Alpha oscillations correlate with the successful inhibition of unattended stimuli. J Cogn Neurosci 23:2494–2502.

Henry MJ, Herrmann B, Kunke D, Obleser J (2017) Aging affects the balance of neural entrainment and top-down neural modulation in the listening brain. Nat Commun 8:15801.

Henry MJ, Obleser J (2012) Frequency modulation entrains slow neural oscillations and optimizes human listening behavior. Proc Natl Acad Sci USA 109:20095–20100.

Hoerl AE, Kennard RW (1970) Ridge Regression: Biased Estimation for Nonorthogonal Problems. Technometrics 12:55–67.

Hong X, Sun J, Bengson JJ, Mangun GR, Tong S (2015) Normal aging selectively diminishes alpha lateralization in visual spatial attention. NeuroImage 106:353–363.

Horton C, D’Zmura M, Srinivasan R (2013) Suppression of competing speech through entrainment of cortical oscillations. J Neurophysiol 109:3082–3093.

Jefferies K, Gale TM (2013) 6-CIT: Six-Item Cognitive Impairment Test. In: Cognitive Screening Instruments, pp 209–218. London: Springer, London.

Jensen O, Mazaheri A (2010) Shaping functional architecture by oscillatory alpha activity: gating by inhibition. Front Hum Neurosci 4: 186.

Kaya EM, Elhilali M (2017) Modelling auditory attention. Philosophical Transactions of the Royal Society B: Biological Sciences 372: 1714.

Kerlin JR, Shahin AJ, Miller LM (2010) Attentional gain control of ongoing cortical speech representations in a “cocktail party.” J Neurosci 30:620–628.

Kimura D (1961) Cerebral dominance and the perception of verbal stimuli. Canadian Journal of Psychology/Revue canadienne de psychologie 15:166–171.

Krakauer JW, Ghazanfar AA, Gomez-Marin A, MacIver MA, Poeppel D (2017) Neuroscience needs behavior: correcting a reductionist bias. Neuron 93:480–490.

Lakatos P, Barczak A, Neymotin SA, McGinnis T, Ross D, Javitt DC, O’Connell MN (2016) Global dynamics of selective attention and its lapses in primary auditory cortex. Nat Neurosci 19:1707–1717.

Lalor EC, Foxe JJ (2010) Neural responses to uninterrupted natural speech can be extracted with precise temporal resolution. Eur J Neurosci 31:189–193.

Leenders MP, Lozano-Soldevilla D, Roberts MJ (2018) Diminished alpha lateralization during working memory but not during attentional cueing in older adults. Cereb Cortex 28:21–32.

Lenth RV (2016) Least-squares means: The R package lsmeans. J Stat Soft 69:1–33.

Luke SG (2017) Evaluating significance in linear mixed-effects models in R. Behav Res 49:1494–1502.

Lüdecke D (2018) Data Visualization for Statistics in Social Science [R package sjPlot version 2.6.1]. Available at: https://CRAN.R-project.org/package=sjPlot.

Melara RD, Rao A, Tong Y (2002) The duality of selection: Excitatory and inhibitory processes in auditory selection attention. Journal of Experimental Psychology: Human Perception and Performance 28:279–306.

Mesgarani N, Chang EF (2012) Selective cortical representation of attended speaker in multi-talker speech perception. Nature 485:233–236.

Mok RM, Myers NE, Wallis G, Nobre AC (2016) Behavioral and neural markers of flexible attention over working memory in aging. Cereb Cortex 26:18311842.

Müller N, Weisz N (2011) Lateralized auditory cortical alpha band activity and interregional connectivity pattern reflect anticipation of target sounds. Cereb Cortex 22:1604–1613.

Nourski KV, Brugge JF (2011) Representation of temporal sound features in the human auditory cortex. Rev Neurosci 22:187–203.

Obleser J, Weisz N (2012) Suppressed Alpha Oscillations Predict Intelligibility of Speech and its Acoustic Details. Cereb Cortex 22:2466–2477.

Oldfield RC (1971) The assessment and analysis of handedness: The Edinburgh inventory. 9:97–113.

O’Sullivan JA, Power AJ, Mesgarani N, Rajaram S, Foxe JJ, Shinn-Cunningham BG, Slaney M, Shamma SA, Lalor EC (2014) Attentional Selection in a Cocktail Party Environment Can Be Decoded from Single-Trial EEG. Cereb Cortex 25:1697–1706.

Peelle JE, Gross J, Davis MH (2013) Phase-Locked Responses to Speech in Human Auditory Cortex are Enhanced During Comprehension. Cereb Cortex 23:1378–1387.

Peirce JW (2007) PsychoPy—Psychophysics software in Python. Journal of Neuroscience Methods 162:8–13.

Posner MI (1980) Orienting of attention. Quarterly Journal of Experimental Psychology 32:3–25.

Power AJ, Foxe JJ, Forde E-J, Reilly RB, Lalor EC (2012) At what time is the cocktail party? A late locus of selective attention to natural speech. Eur J Neurosci 35:1497–1503.

Presacco A, Simon JZ, Anderson S (2016a) Effect of informational content of noise on speech representation in the aging midbrain and cortex. J Neurophysiol 116:2356–2367.

Presacco A, Simon JZ, Anderson S (2016b) Evidence of degraded representation of speech in noise, in the aging midbrain and cortex. J Neurophysiol 116:2346–2355.

R Core Team (2018) R: A language and environment for statistical computing. Available at: https://www.R-project.org/.

Rihs TA, Michel CM, Thut G (2007) Mechanisms of selective inhibition in visual spatial attention are indexed by α-band EEG synchronization. Eur J Neurosci 25:603–610.

Sadaghiani S, Kleinschmidt A (2016) Brain Networks and α-Oscillations: Structural and Functional Foundations of Cognitive Control. Trends Cogn Sci 20:805–817.

Sander MC, Werkle-Bergner M, Lindenberger U (2012) Amplitude modulations and inter-trial phase stability of alpha-oscillations differentially reflect working memory constraints across the lifespan. NeuroImage 59:646–654.

Schroeder CE, Lakatos P (2009) Low-frequency neuronal oscillations as instruments of sensory selection. Trends Neurosci 32:9–18.

Schroeder CE, Wilson DA, Radman T, Scharfman H, Lakatos P (2010) Dynamics of Active Sensing and perceptual selection. Current Opinion in Neurobiology 20:172–176.

Shinn-Cunningham BG (2008) Object-based auditory and visual attention. Trends Cogn Sci 12:182–186.

Shinn-Cunningham BG, Best V (2008) Selective Attention in Normal and Impaired Hearing. Trends in Amplification 12:283–299.

Smulders FTY, Oever ten S, Donkers FCL, Quaedflieg CWEM, van de Ven V (2018) Single-trial log transformation is optimal in frequency analysis of resting EEG alpha. Eur J Neurosci 44:94–14.

Sohoglu E, Peelle JE, Carlyon RP, Davis MH (2012) Predictive Top-Down Integration of Prior Knowledge during Speech Perception. J Neurosci 32:8443–8453.

Tune S, Wöstmann M, Obleser J (2018) Probing the limits of alpha power lateralisation as a neural marker of selective attention in middle-aged and older listeners. Eur J Neurosci 25:1926–14.

van Ede F, Köster M, Maris E (2012) Beyond establishing involvement: quantifying the contribution of anticipatory α- and β-band suppression to perceptual improvement with attention. J Neurophysiol 108:2352–2362.

Wöstmann M, Herrmann B, Maess B, Obleser J (2016) Spatiotemporal dynamics of auditory attention synchronize with speech. Proc Natl Acad Sci USA 113:3873–3878.

Wöstmann M, Lim S-J, Obleser J (2017) The human neural alpha response to speech is a proxy of attentional control. Cereb Cortex 27:3307–3317.

Wöstmann M, Obleser J (2016) Acoustic detail but not predictability of task-irrelevant speech disrupts working memory. Front Hum Neurosci 10:201–209.

Wöstmann M, Vosskuhl J, Obleser J, Herrmann CS (2018) Opposite effects of lateralised transcranial alpha versus gamma stimulation on auditory spatial attention. Brain Stimulation: 11:752–758.

Zion Golumbic EMZ, Ding N, Bickel S, Lakatos P, Schevon CA, McKhann GM, Goodman RR, Emerson R, Mehta AD, Simon JZ, Poeppel D, Schroeder CE (2013) Mechanisms Underlying Selective Neuronal Tracking of Attended Speech at a “Cocktail Party”. Neuron 77:980–991.

