## Supplemental Information for "Individual listening success explained by synergistic interaction of two distinct neural filters"

**Supplemental Information**
**for**

### Individual listening success explained by synergistic interaction of two distinct neural filters

Sarah Tune*, Lorenz Fiedler, Mohsen Alavash, Jonas Obleser*

Department of Psychology, University of Lübeck, 23562 Lübeck, Germany

** Author correspondence*:
Sarah Tune, Jonas Obleser
Department of Psychology
University of Lübeck
Maria-Goeppert-Str. 9a
23562 Lübeck


#### Supplemental figures

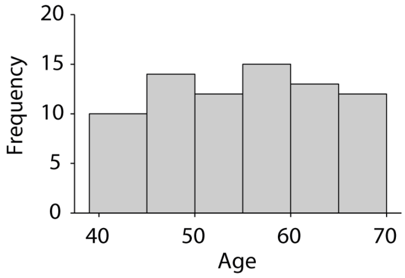


**Figure 2–supplement 1. Histogram showing age distribution of N = 76 participants across six age bins.**

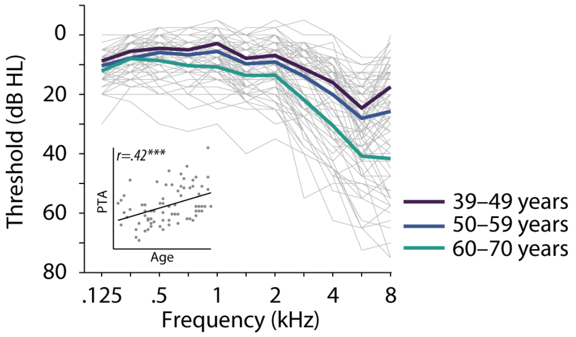


**Figure 2–supplement 2. Individual and mean air conduction thresholds averaged across the left and right ear.** Thin grey lines show air conduction pure-tone thresholds for individual participants, thick colored lines indicate average thresholds grouped across three age bins. Inset scatterplot shows positive Pearson correlation (r = .42, p < .001) of pure-tone average (PTA; mean across left and right ear at 500, 1000, 2000 and 4000 Hz) and age. Grey dots show thresholds for individual participants.

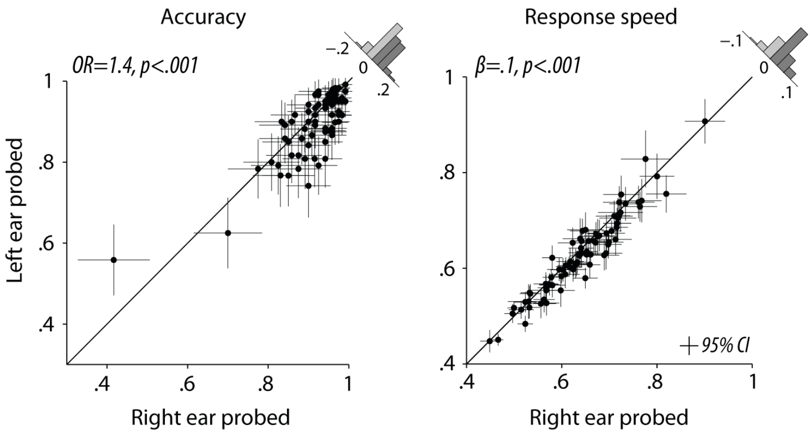


**Figure 2–supplement 3. Right-ear advantage for accuracy and response speed.** Individual right-ear advantage for N=76 participants displayed for accuracy and response speed, respectively. Black dots indicate individual trial-averages ± bootstrapped 95% confidence intervals. Histograms show the distribution of the difference of right-ear vs. left-ear probed trials across all participants. OR: Odds ratio parameter estimate from generalized linear mixed-effects models; *β:* slope parameter estimate from general linear mixed-effects models.

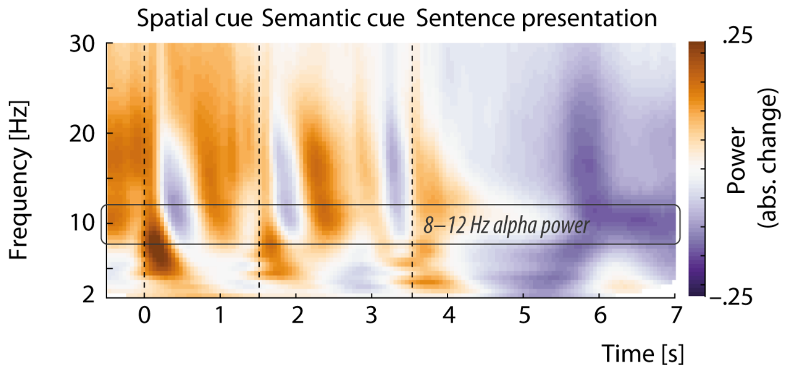


**Figure 3–supplement 1. Whole-trial overall oscillatory power averaged across all trials, electrodes, and N = 76 participants.** Vertical dotted lines indicate the onset of the two visual listening cues and the presentation of concurrent speech, respectively. Power changes are calculated as absolute change relative to a whole-trial (0–7 s) baseline interval.

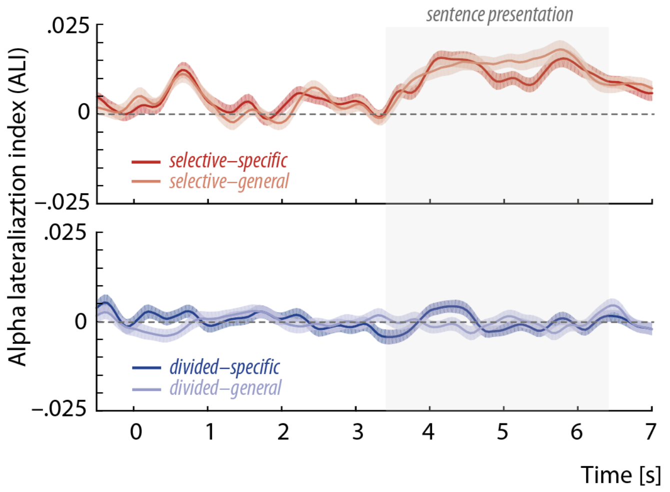


**Figure 3–supplement 2. Grand-average whole-trial alpha lateralization index (ALI) calculated without adjustment for overall differences in power across hemispheres.** Top row (red traces) shows ALI values (± 95% confidence intervals) for selective-attention trials separately per semantic cue; bottom row (blue traces) shows ALI values for divided-attention trials. Grey intervals marks interval of dichotic sentence presentation.

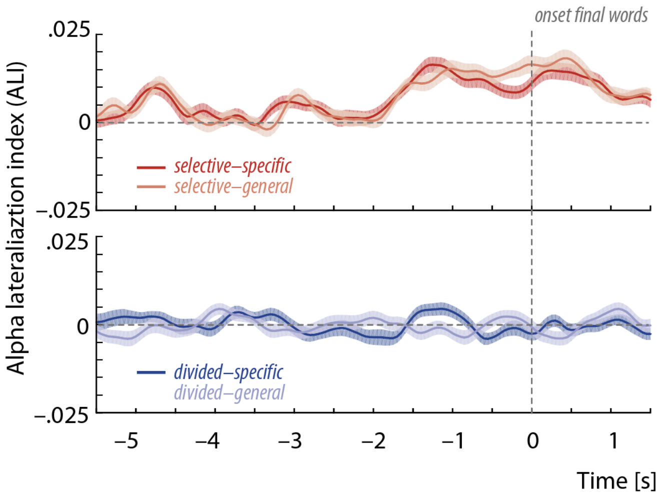


**Figure 3–supplement 3. Grand-average whole-trial alpha lateralization index (ALI) time-locked to the onset of the sentence-final target words.** Top row (red traces) shows ALI values (± 95% confidence intervals) for selective-attention trials separately per semantic cue; bottom row (blue traces) shows ALI values for divided-attention trials. Vertical dotted lines marks onset of time-locked sentence-final word presentation.

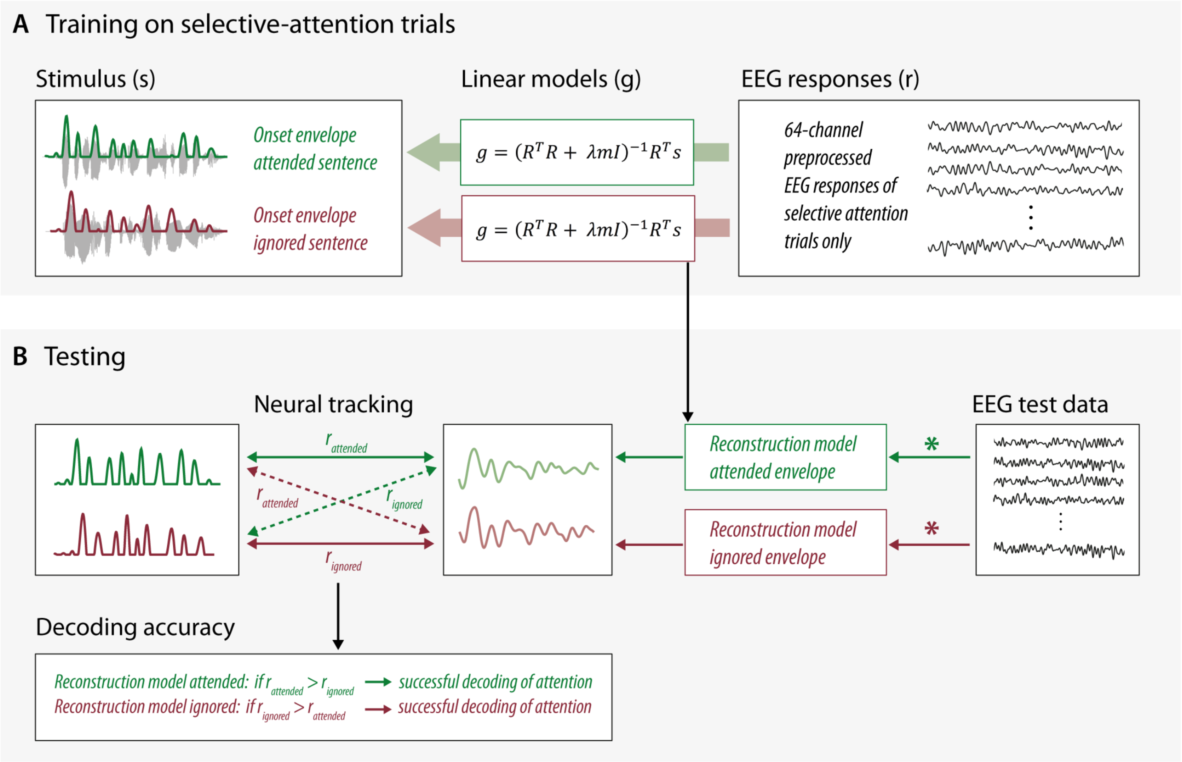


**Figure 4–supplement 1. Training and testing of envelope reconstruction models.**
(A) Linear backward models were estimated using single-trial onset envelopes and preprocessed EEG responses of selective attention trials, only. Single-subject models for the attended and ignored speech stream were trained separately.
(B) Following a leave-one-out procedure for selective-attention trials, single-trial envelope reconstructions were computed by convolving the EEG signal of the test trial with the trained reconstructions models averaged across all but the tested trial. For the reconstruction of envelope presented in divided-attention trials, we used the average of all single-trial decoder models estimated in training. Neural tracking strength was quantified as the Pearson correlation coefficient of the reconstructed and presented envelopes yielding four coefficients per trial and subject. Decoding accuracy was assessed separately for the reconstruction model of the attended and ignored envelope, respectively. For the former, a participant's attention was correctly decoded if the correlation coefficient for the comparison of the reconstructed with the attended envelope ($r_{attended}$) was larger than that of the comparison with the ignored envelope ($r_{ignored}$). The opposite relationship holds for the decoding accuracy of the ignored envelope reconstruction model.

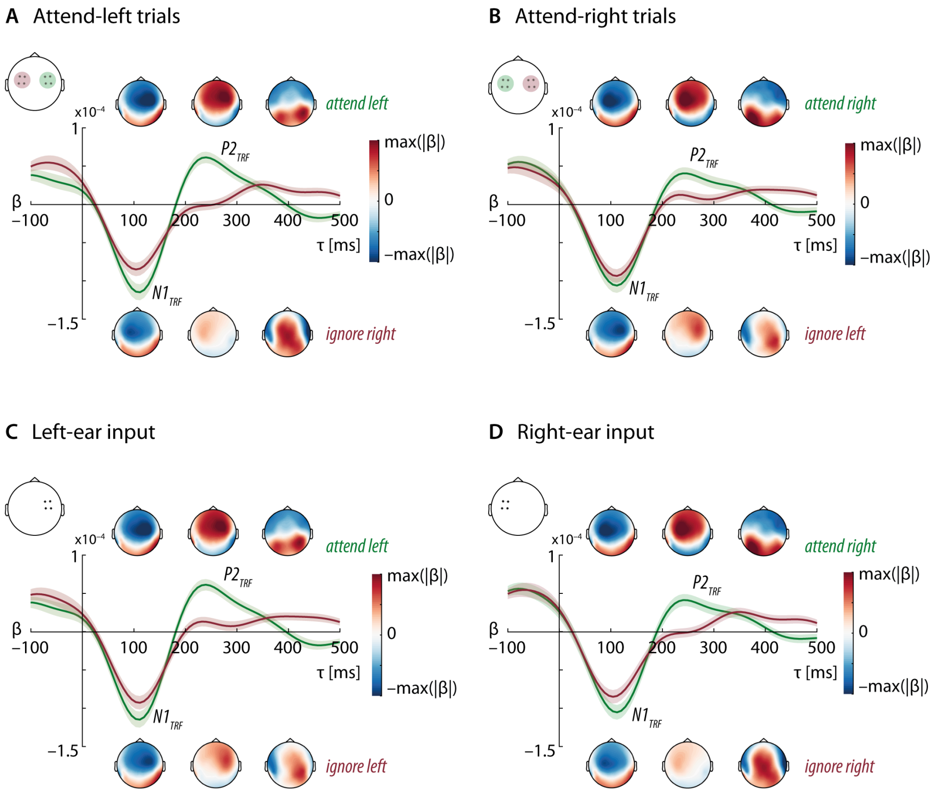


**Figure 4–supplement 2. Encoding of attended and ignored speech shown separately per probed-ear condition.** Topographical distribution of linear backward models weight shown in line with forward-transformed temporal response functions (TRFs) averaged across all *N*=76 participants. TRFs are averaged across four fronto-central electrodes in the left (FC3, FC5, C3, C5) or right hemisphere (FC4, FC6, C4, C6) as indicated by the topographical maps displayed on the top left side of each panel. Error bands indicate 95% confidence intervals. Backward model weights are averaged for the time lag intervals of 90–110 ms, 200–250 ms, and 350–450 ms, respectively.
(A, B) Model results shown for attend-left/ignore-right and attend-right/ignore-left trials, respectively.
(C, D) Model results shown per stimulus presentation side. To-be-attended and to-be-ignored stimuli were presented in different trials.

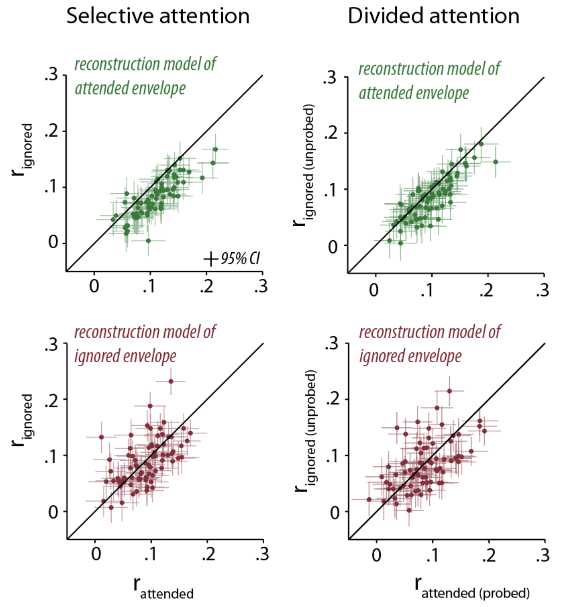


**Figure 4–supplement 3. Neural tracking strength in selective and divided-attention trials.** Single-subject neural tracking strength compared for attended and ignored speech. Top row shows results of the reconstruction model for the attended envelope; bottom row shows results of the reconstruction model for the ignored envelope. Error bars index 95% bootstrapped confidence intervals.

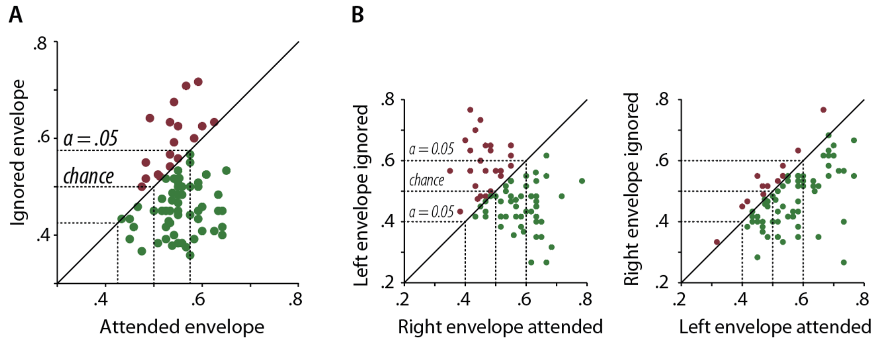


**Figure 4–supplement 4. Single-subject decoding accuracy of the attended and ignored reconstruction models in selective-attention trials.** (A) Single-subject decoding accuracy for attended and ignored speech when trials are pooled across probed-ear conditions. (B) Single-subject decoding accuracy shown separately per attend-right/ignore-left (left plot) and attend-left/ignore-right trials (right plot).

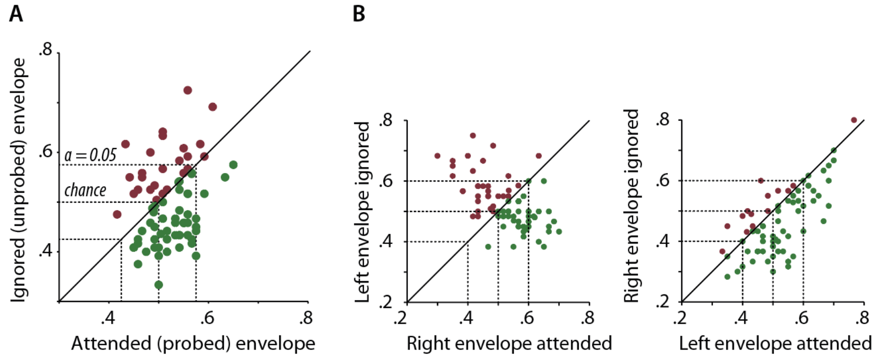


**Figure 4–supplement 5. Single-subject decoding accuracy of the attended and ignored reconstruction models in divided-attention trials.** (A) Single-subject decoding accuracy for attended and ignored speech when trials are pooled across probed-ear conditions. (B) Single-subject decoding accuracy shown separately per attend-right/ignore-left (left plot) and attend-left/ignore-right trials (right plot).

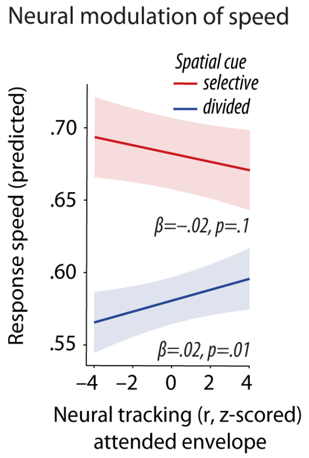


**Figure 5–supplement 1. Modulation of response speed by neural tracking strength ($r_{attended}$) under each attention condition.** Fixed-effect relationship of attended-speech tracking on response speed as predicted by the corresponding brain-behavior linear mixed-effects model (for details see Table S2). Error bands index 95% confidence intervals. Under divided attention, stronger neural tracking of attended speech leads to faster responses.

### Statistical model details

#### Table S1: Brain-behavior model predicting accuracy

| | Accuracy | | | | | Accuracy:Attend-L trials | | | | | Accuracy:Attend-R trials | | | | |
|---|---|---|---|---|---|---|---|---|---|---|---|---|---|---|---|
| *Predictors* | *Odds Ratios* | *std. Error* | *CI* | *z-value* | *p* | *Odds Ratios* | *std. Error* | *CI* | *z-value* | *p* | *Odds Ratios* | *std. Error* | *CI* | *z-value* | *p* |
| Intercept | 24.06 | 0.12 | 18.97 – 30.52 | 26.20 | **<0.001** | 20.95 | 0.13 | 16.15 – 27.17 | 22.90 | **<0.001** | 29.14 | 0.14 | 21.98 – 38.62 | 23.45 | **<0.001** |
| Spatial cue (selective) | 3.37 | 0.14 | 2.56 – 4.44 | 8.65 | **<0.001** | 3.15 | 0.16 | 2.31 – 4.28 | 7.29 | **<0.001** | 3.67 | 0.19 | 2.52 – 5.35 | 6.77 | **<0.001** |
| Semantic cue (specific) | 1.02 | 0.13 | 0.80 – 1.31 | 0.16 | 0.876 | 0.93 | 0.14 | 0.70 – 1.23 | -0.51 | 0.610 | 1.16 | 0.15 | 0.87 – 1.55 | 1.00 | 0.319 |
| Probed ear (right) | 1.38 | 0.09 | 1.15 – 1.66 | 3.50 | **<0.001** | | | | | | | | | | |
| First sentence (to-be-attended) | 0.99 | 0.06 | 0.88 – 1.12 | -0.09 | 0.931 | 0.94 | 0.09 | 0.80 – 1.12 | -0.66 | 0.510 | 1.01 | 0.10 | 0.84 – 1.23 | 0.14 | 0.891 |
| Age (z-scored) | 0.89 | 0.11 | 0.71 – 1.11 | -1.06 | 0.291 | 0.88 | 0.12 | 0.69 – 1.13 | -0.98 | 0.326 | 0.87 | 0.13 | 0.68 – 1.13 | -1.05 | 0.294 |
| PTA (z-scored) | 0.75 | 0.11 | 0.61 – 0.92 | -2.72 | **0.007** | 0.74 | 0.12 | 0.59 – 0.94 | -2.51 | **0.012** | 0.76 | 0.12 | 0.60 – 0.96 | -2.30 | **0.021** |
| Neural tracking attended (z-scored) | 0.99 | 0.04 | 0.93 – 1.06 | -0.22 | 0.824 | 0.91 | 0.05 | 0.82 – 1.01 | -1.82 | 0.068 | 1.06 | 0.05 | 0.96 – 1.16 | 1.12 | 0.264 |
| Neural tracking ignored (z-scored) | 1.04 | 0.03 | 0.97 – 1.11 | 1.04 | 0.299 | 1.12 | 0.05 | 1.01 – 1.24 | 2.20 | **0.028** | 0.96 | 0.05 | 0.88 – 1.06 | -0.76 | 0.450 |
| ALI (z-scored) | 1.00 | 0.03 | 0.94 – 1.07 | 0.15 | 0.880 | 0.99 | 0.04 | 0.91 – 1.08 | -0.29 | 0.772 | 1.02 | 0.05 | 0.93 – 1.13 | 0.49 | 0.626 |
| Probed ear x Semantic cue | 1.17 | 0.12 | 0.92 – 1.48 | 1.29 | 0.196 | | | | | | | | | | |
| First sentence x Semantic Cue | 2.19 | 0.12 | 1.73 – 2.77 | 6.47 | **<0.001** | 2.40 | 0.17 | 1.72 – 3.33 | 5.21 | **<0.001** | 1.87 | 0.19 | 1.30 – 2.70 | 3.34 | **0.001** |
| Age x Spatial cue | 1.15 | 0.08 | 0.99 – 1.33 | 1.79 | 0.074 | 1.13 | 0.09 | 0.95 – 1.34 | 1.33 | 0.185 | 1.14 | 0.12 | 0.90 – 1.46 | 1.08 | 0.281 |
| Probed ear x Neural tracking attended | 1.18 | 0.07 | 1.03 – 1.35 | 2.43 | **0.015** | | | | | | | | | | |
| Probed ear x Neural tracking ignored | 0.85 | 0.07 | 0.75 – 0.98 | -2.28 | **0.023** | | | | | | | | | | |
| Probed ear x ALI | 1.02 | 0.06 | 0.89 – 1.15 | 0.25 | 0.805 | | | | | | | | | | |
| Neural tracking attended x ALI | 0.99 | 0.04 | 0.93 – 1.06 | -0.19 | 0.850 | 0.94 | 0.05 | 0.85 – 1.05 | -1.07 | 0.287 | 1.05 | 0.05 | 0.96 – 1.16 | 1.13 | 0.258 |
| Neural tracking ignored x ALI | 1.09 | 0.03 | 1.02 – 1.17 | 2.48 | **0.013** | 1.17 | 0.05 | 1.05 – 1.29 | 2.97 | **0.003** | 1.00 | 0.05 | 0.91 – 1.09 | -0.03 | 0.976 |
| Probed ear x Neural tracking attended x ALI | 1.11 | 0.07 | 0.97 – 1.28 | 1.52 | 0.128 | | | | | | | | | | |
| Probed ear x Neural tracking ignored x ALI | 0.86 | 0.07 | 0.75 – 0.98 | -2.22 | **0.026** | | | | | | | | | | |
| **Random Effects** | | | | | | | | | | | | | | | |
| $\sigma^2$ | 3.29 | | | | | 3.29 | | | | | 3.29 | | | | |
| $\tau_{00}$ | 0.66 $_{pair}$ | | | | | 0.72 $_{pair}$ | | | | | 0.67 $_{pair}$ | | | | |
| | 0.73 $_{subj}$ | | | | | 0.81 $_{subj}$ | | | | | 0.84 $_{subj}$ | | | | |
| $\tau_{11}$ | 0.10 $_{subj.spatialSelective}$ | | | | | 0.02 $_{subj.spatialSelective}$ | | | | | 0.28 $_{subj.spatialSelective}$ | | | | |
| | 0.26 $_{subj.probedRight}$ | | | | | | | | | | | | | | |
| $\rho_{01}$ | 0.64 $_{subj.spatialSelective}$ | | | | | 1.00 $_{subj}$ | | | | | 0.53 $_{subj}$ | | | | |
| | 0.08 $_{subj.probedRight}$ | | | | | | | | | | | | | | |
| Observations | 17532 | | | | | 8759 | | | | | 8773 | | | | |
| Marginal $R^2$ / Conditional $R^2$ | 0.109 / 0.387 | | | | | 0.100 / 0.386 | | | | | 0.110 / 0.399 | | | | |

#### Table S2: Brain-behavior model predicting response speed

| | Speed (z-scored) | | | | | Speed: Selective attention trials | | | | | Speed: Divided attention trials | | | | |
|---|---|---|---|---|---|---|---|---|---|---|---|---|---|---|---|
| *Predictors* | *Estimates* | *std. Error* | *CI* | *t-value* | *p* | *Estimates* | *std. Error* | *CI* | *t-value* | *p* | *Estimates* | *std. Error* | *CI* | *t-value* | *p* |
| Intercept | -0.034 | 0.055 | -0.141 – 0.074 | -0.616 | 0.538 | 0.255 | 0.073 | 0.113 – 0.398 | 3.516 | **<0.001** | -0.321 | 0.046 | -0.410 – -0.231 | -7.019 | **<0.001** |
| Spatial cue (selective) | 0.575 | 0.050 | 0.477 – 0.672 | 11.555 | **<0.001** | | | | | | | | | | |
| Semantic cue (specific) | 0.192 | 0.035 | 0.122 – 0.261 | 5.416 | **<0.001** | 0.243 | 0.054 | 0.136 – 0.349 | 4.472 | **<0.001** | 0.140 | 0.043 | 0.055 – 0.225 | 3.234 | **0.001** |
| Probed ear (right) | 0.063 | 0.015 | 0.033 – 0.093 | 4.101 | **<0.001** | 0.049 | 0.024 | 0.002 – 0.095 | 2.065 | **0.039** | 0.080 | 0.018 | 0.044 – 0.116 | 4.330 | **<0.001** |
| First sentence (to-be-attended) | 0.008 | 0.013 | -0.017 – 0.033 | 0.646 | 0.518 | 0.009 | 0.019 | -0.028 – 0.047 | 0.491 | 0.623 | 0.003 | 0.016 | -0.029 – 0.035 | 0.186 | 0.852 |
| Age (z-scored) | -0.118 | 0.040 | -0.196 – -0.041 | -2.988 | **0.003** | -0.122 | 0.072 | -0.263 – 0.018 | -1.709 | 0.088 | -0.120 | 0.045 | -0.207 – -0.032 | -2.676 | **0.007** |
| PTA (z-scored) | -0.034 | 0.039 | -0.111 – 0.043 | -0.864 | 0.387 | -0.060 | 0.071 | -0.198 – 0.078 | -0.847 | 0.397 | -0.034 | 0.044 | -0.120 – 0.053 | -0.765 | 0.444 |
| Neural tracking attended (z-scored) | 0.003 | 0.007 | -0.010 – 0.016 | 0.446 | 0.655 | -0.017 | 0.010 | -0.037 – 0.004 | -1.608 | 0.108 | 0.022 | 0.009 | 0.005 – 0.040 | 2.540 | **0.011** |
| Neural tracking ignored (z-scored) | 0.001 | 0.007 | -0.012 – 0.014 | 0.182 | 0.856 | 0.005 | 0.010 | -0.015 – 0.025 | 0.465 | 0.642 | -0.003 | 0.009 | -0.020 – 0.014 | -0.376 | 0.707 |
| ALI (z-scored) | 0.009 | 0.006 | -0.003 – 0.021 | 1.417 | 0.156 | 0.005 | 0.010 | -0.014 – 0.024 | 0.524 | 0.601 | 0.015 | 0.008 | -0.001 – 0.031 | 1.808 | 0.071 |
| Spatial cue x Semantic cue | 0.103 | 0.068 | -0.030 – 0.236 | 1.520 | 0.129 | | | | | | | | | | |
| ALI x Neural tracking attended | 0.006 | 0.007 | -0.007 – 0.019 | 0.945 | 0.344 | 0.005 | 0.010 | -0.014 – 0.025 | 0.533 | 0.594 | 0.008 | 0.008 | -0.008 – 0.025 | 0.983 | 0.326 |
| ALI x Neural tracking ignored | 0.001 | 0.007 | -0.012 – 0.014 | 0.198 | 0.843 | 0.003 | 0.010 | -0.016 – 0.023 | 0.343 | 0.731 | -0.002 | 0.008 | -0.018 – 0.014 | -0.217 | 0.828 |
| Neural tracking attended x Spatial Cue | -0.039 | 0.013 | -0.065 – -0.013 | -2.891 | **0.004** | | | | | | | | | | |
| Neural tracking ignored x Spatial Cue | 0.007 | 0.013 | -0.019 – 0.034 | 0.544 | 0.587 | | | | | | | | | | |
| Neural tracking attended x First sentence | 0.024 | 0.014 | -0.002 – 0.051 | 1.777 | 0.076 | 0.021 | 0.021 | -0.019 – 0.061 | 1.014 | 0.311 | 0.028 | 0.017 | -0.006 – 0.062 | 1.603 | 0.109 |
| Neural tracking ignored x First sentence | -0.014 | 0.013 | -0.041 – 0.012 | -1.068 | 0.286 | -0.010 | 0.020 | -0.050 – 0.030 | -0.507 | 0.612 | -0.018 | 0.017 | -0.052 – 0.015 | -1.072 | 0.284 |
| **Random Effects** | | | | | | | | | | | | | | | |
| $\sigma^2$ | 0.60 | | | | | 0.72 | | | | | 0.46 | | | | |
| $\tau_{00}$ | 0.06 $_{pair}$ | | | | | 0.07 $_{pair}$ | | | | | 0.05 $_{pair}$ | | | | |
| | 0.21 $_{subj}$ | | | | | 0.35 $_{subj}$ | | | | | 0.12 $_{subj}$ | | | | |
| $\tau_{11}$ | 0.10 $_{subj.spatialSelective}$ | | | | | 0.01 $_{subj.semanticSpecific}$ | | | | | 0.01 $_{subj.semanticSpecific}$ | | | | |
| | 0.01 $_{subj.semanticSpecific}$ | | | | | 0.02 $_{subj.probedRight}$ | | | | | 0.01 $_{subj.probedRight}$ | | | | |
| | 0.01 $_{subj.probedRight}$ | | | | | | | | | | | | | | |
| $\rho_{01}$ | 0.77 $_{subj.spatialSelective}$ | | | | | 0.51 $_{subj.semanticSpecific}$ | | | | | -0.19 $_{subj.semanticSpecific}$ | | | | |
| | 0.30 $_{subj.semanticSpecific}$ | | | | | -0.15 $_{subj.probedRight}$ | | | | | 0.28 $_{subj.probedRight}$ | | | | |
| | 0.09 $_{subj.probedRight}$ | | | | | | | | | | | | | | |
| Observations | 16128 | | | | | 8542 | | | | | 7586 | | | | |
| Marginal $R^2$ / Conditional $R^2$ | 0.110 / 0.407 | | | | | 0.034 / 0.394 | | | | | 0.040 / 0.301 | | | | |

#### Table S3: Cue-driven neural modulation model predicting alpha power lateralization during sentence presentation

| | Alpha power lateralization (sentences) | | | | |
|---|---|---|---|---|---|
| *Predictors* | *Estimates* | *std. Error* | *CI* | *t-value* | *p* |
| Intercept | -0.000 | 0.012 | -0.024 – 0.023 | -0.034 | 0.973 |
| Spatial cue (selective) | 0.214 | 0.022 | 0.171 – 0.257 | 9.743 | **<0.001** |
| Probed ear (right) | 0.031 | 0.070 | -0.106 – 0.168 | 0.448 | 0.654 |
| Age (z-scored) | -0.010 | 0.012 | -0.034 – 0.013 | -0.862 | 0.389 |
| Age x Spatial Cue | -0.025 | 0.022 | -0.067 – 0.017 | -1.159 | 0.247 |
| **Random Effects** | | | | | |
| $\sigma^2$ | 0.89 | | | | |
| $\tau_{00\ subj}$ | 0.01 | | | | |
| $\tau_{11\ subj.spatialSelective}$ | 0.02 | | | | |
| $\tau_{11\ subj.probedRight}$ | 0.36 | | | | |
| $\rho_{01\ subj.spatialSelective}$ | 0.99 | | | | |
| $\rho_{01\ subj.probedRight}$ | 0.08 | | | | |
| Observations | 17824 | | | | |
| Marginal $R^2$ / Conditional $R^2$ | 0.012 / 0.113 | | | | |

#### Table S4: Cue-driven neural modulation model predicting alpha power lateralization during spatial-cue presentation

| | Alpha power lateralization (spatial cue) | | | | |
|---|---|---|---|---|---|
| *Predictors* | *Estimates* | *std. Error* | *CI* | *t-value* | *p* |
| Intercept | -0.000 | 0.008 | -0.016 – 0.016 | -0.015 | 0.988 |
| Spatial cue (selective) | 0.076 | 0.015 | 0.046 – 0.105 | 5.065 | **<0.001** |
| Probed ear (right) | -0.050 | 0.015 | -0.080 – -0.021 | -3.355 | **0.001** |
| Age (z-scored) | 0.009 | 0.008 | -0.008 – 0.025 | 1.027 | 0.304 |
| Age x Spatial Cue | -0.030 | 0.015 | -0.059 – -0.001 | -2.011 | **0.044** |
| **Random Effects** | | | | | |
| $\sigma^2$ | 1.00 | | | | |
| $\tau_{00\ subj}$ | 0.00 | | | | |
| Observations | 17824 | | | | |
| Marginal $R^2$ / Conditional $R^2$ | 0.002 / 0.003 | | | | |

#### Table S5: Cue-driven neural modulation model predicting neural tracking of attended envelope

| | Neural tracking attended envelope (r-attended) | | | | |
|---|---|---|---|---|---|
| *Predictors* | *Estimates* | *std. Error* | *CI* | *t-value* | *p* |
| Intercept | -0.005 | 0.027 | -0.058 – 0.049 | -0.165 | 0.869 |
| Spatial cue (selective) | 0.079 | 0.026 | 0.028 – 0.129 | 3.073 | **0.002** |
| Semantic cue (specific) | 0.019 | 0.026 | -0.031 – 0.069 | 0.753 | 0.452 |
| Probed ear (right) | -0.041 | 0.024 | -0.087 – 0.005 | -1.736 | 0.083 |
| First sentence (to-be-attended) | 0.338 | 0.015 | 0.309 – 0.367 | 23.034 | **<0.001** |
| Age (z-scored) | 0.057 | 0.025 | 0.008 – 0.105 | 2.286 | **0.022** |
| **Random Effects** | | | | | |
| $\sigma^2$ | 0.89 | | | | |
| $\tau_{00\ pair}$ | 0.03 | | | | |
| $\tau_{00\ subj}$ | 0.04 | | | | |
| $\tau_{11\ subj.probedRight}$ | 0.03 | | | | |
| $\rho_{01\ subj}$ | -0.21 | | | | |
| Observations | 17533 | | | | |
| Marginal $R^2$ / Conditional $R^2$ | 0.033 / 0.111 | | | | |

#### Table S6: Cue-driven neural modulation model predicting neural tracking of ignored envelope

| | Neural tracking ignored envelope (r-ignored) | | | | | Neural tracking: selective attention | | | | | Neural tracking: divided attention | | | | |
|---|---|---|---|---|---|---|---|---|---|---|---|---|---|---|---|
| *Predictors* | *Estimates* | *std. Error* | *CI* | *t-value* | *p* | *Estimates* | *std. Error* | *CI* | *t-value* | *p* | *Estimates* | *std. Error* | *CI* | *t-value* | *p* |
| Intercept | -0.004 | 0.030 | -0.064 – 0.056 | -0.132 | 0.895 | 0.015 | 0.032 | -0.038 – 0.067 | 0.458 | 0.648 | -0.023 | 0.034 | -0.078 – 0.033 | -0.667 | 0.506 |
| Spatial cue (selective) | 0.037 | 0.024 | -0.010 – 0.085 | 1.537 | 0.124 | | | | | | | | | | |
| Semantic cue (specific) | -0.025 | 0.024 | -0.072 – 0.023 | -1.014 | 0.311 | -0.084 | 0.033 | -0.138 – -0.030 | -2.560 | **0.012** | 0.033 | 0.036 | -0.026 – 0.092 | 0.927 | 0.356 |
| Probed ear (right) | 0.077 | 0.022 | 0.034 – 0.120 | 3.530 | **<0.001** | 0.041 | 0.027 | -0.003 – 0.086 | 1.533 | 0.130 | 0.126 | 0.028 | 0.081 – 0.172 | 4.580 | **<0.001** |
| First sentence (to-be-attended) | -0.007 | 0.021 | -0.048 – 0.033 | -0.360 | 0.719 | -0.009 | 0.026 | -0.052 – 0.033 | -0.371 | 0.711 | -0.009 | 0.024 | -0.048 – 0.030 | -0.375 | 0.709 |
| Age (z-scored) | 0.021 | 0.029 | -0.036 – 0.077 | 0.714 | 0.475 | 0.032 | 0.029 | -0.015 – 0.079 | 1.110 | 0.271 | 0.011 | 0.030 | -0.038 – 0.061 | 0.372 | 0.711 |
| Spatial Cue x Semantic cue | -0.116 | 0.048 | -0.211 – -0.021 | -2.397 | **0.017** | | | | | | | | | | |
| Spatial Cue x Probed ear | -0.079 | 0.029 | -0.135 – -0.022 | -2.734 | **0.006** | | | | | | | | | | |
| Semantic Cue x Probed ear | 0.049 | 0.029 | -0.008 – 0.105 | 1.676 | 0.094 | 0.108 | 0.040 | 0.042 – 0.174 | 2.688 | **0.007** | -0.002 | 0.042 | -0.070 – 0.067 | -0.044 | 0.965 |
| **Random Effects** | | | | | | | | | | | | | | | |
| $\sigma^2$ | 0.91 | | | | | 0.89 | | | | | 0.92 | | | | |
| $\tau_{00}$ | 0.02 $_{pair}$ | | | | | 0.02 $_{pair}$ | | | | | 0.02 $_{pair}$ | | | | |
| | 0.06 $_{subj}$ | | | | | 0.06 $_{subj}$ | | | | | 0.06 $_{subj}$ | | | | |
| $\tau_{11}$ | 0.02 $_{subj.probedRight}$ | | | | | 0.02 $_{subj.probedRight}$ | | | | | 0.02 $_{subj.probedRight}$ | | | | |
| | 0.02 $_{subj.att\_firstYes}$ | | | | | 0.02 $_{subj.att\_firstYes}$ | | | | | 0.01 $_{subj.att\_firstYes}$ | | | | |
| $\rho_{01}$ | 0.07 $_{subj.probedRight}$ | | | | | 0.03 $_{subj.probedRight}$ | | | | | 0.18 $_{subj.probedRight}$ | | | | |
| | 0.07 $_{subj.att\_firstYes}$ | | | | | 0.29 $_{subj.att\_firstYes}$ | | | | | -0.02 $_{subj.att\_firstYes}$ | | | | |
| Observations | 17533 | | | | | 8980 | | | | | 8553 | | | | |
| Marginal $R^2$ / Conditional $R^2$ | 0.004 / 0.094 | | | | | 0.004 / 0.093 | | | | | 0.004 / 0.097 | | | | |

#### Table S7: Neural covariation model predicting neural tracking of the attended envelope from alpha power

| | Neural tracking attended envelope (r-attended) | | | | |
|---|---|---|---|---|---|
| *Predictors* | *Estimates* | *std. Error* | *CI* | *t-value* | *p* |
| Intercept | 0.000 | 0.026 | -0.042 – 0.043 | 0.009 | 0.993 |
| Alpha power ipsi | -0.013 | 0.014 | -0.036 – 0.010 | -0.917 | 0.359 |
| Alpha power contra | 0.014 | 0.014 | -0.009 – 0.036 | 0.986 | 0.324 |
| Alpha power ipsi x Alpha power contra | -0.001 | 0.002 | -0.005 – 0.002 | -0.621 | 0.535 |
| **Random Effects** | | | | | |
| $\sigma^2$ | 0.95 | | | | |
| $\tau_{00\ subj}$ | 0.05 | | | | |
| Observations | 17824 | | | | |
| Marginal $R^2$ / Conditional $R^2$ | 0.000 / 0.046 | | | | |

#### Table S8: Neural covariation model predicting neural tracking of the ignored envelope from alpha power

| | Neural tracking ignored envelope (r-ignored) | | | | |
|---|---|---|---|---|---|
| *Predictors* | *Estimates* | *std. Error* | *CI* | *t-value* | *p* |
| Intercept | -0.002 | 0.029 | -0.050 – 0.046 | -0.070 | 0.944 |
| Alpha power ipsi | -0.005 | 0.014 | -0.028 – 0.018 | -0.368 | 0.713 |
| Alpha power contra | 0.002 | 0.014 | -0.020 – 0.024 | 0.147 | 0.883 |
| Alpha power ipsi x Alpha power contra | 0.001 | 0.002 | -0.002 – 0.004 | 0.465 | 0.642 |
| **Random Effects** | | | | | |
| $\sigma^2$ | 0.94 | | | | |
| $\tau_{00\ subj}$ | 0.06 | | | | |
| Observations | 17824 | | | | |
| Marginal $R^2$ / Conditional $R^2$ | 0.000 / 0.060 | | | | |

#### Table 9: Neural covariation model predicting neural tracking of the attended and ignored envelope from alpha power lateralization (ALI)

| | Neural tracking attended envelope (r-attended) | | | | | Neural tracking ignored envelope (r-ignored) | | | | |
|---|---|---|---|---|---|---|---|---|---|---|
| *Predictors* | *Estimates* | *std. Error* | *CI* | *t-value* | *p* | *Estimates* | *std. Error* | *CI* | *t-value* | *p* |
| Intercept | -0.001 | 0.025 | -0.042 – 0.041 | -0.030 | 0.976 | -0.002 | 0.029 | -0.049 – 0.046 | -0.054 | 0.957 |
| ALI (z-scored) | -0.015 | 0.009 | -0.031 – 0.000 | -1.627 | 0.112 | -0.000 | 0.010 | -0.016 – 0.016 | -0.006 | 0.995 |
| Spatial cue (selective) | 0.075 | 0.015 | 0.051 – 0.099 | 5.105 | **<0.001** | 0.034 | 0.015 | 0.010 – 0.058 | 2.308 | **0.021** |
| Semantic cue (specific) | 0.012 | 0.015 | -0.012 – 0.036 | 0.844 | 0.399 | -0.029 | 0.015 | -0.053 – -0.005 | -2.006 | **0.045** |
| Probed ear (right) | -0.019 | 0.015 | -0.043 – 0.006 | -1.255 | 0.209 | 0.088 | 0.015 | 0.064 – 0.112 | 5.970 | **<0.001** |
| Age (z-scored) | 0.052 | 0.025 | 0.010 – 0.093 | 2.056 | **0.043** | 0.022 | 0.029 | -0.026 – 0.069 | 0.753 | 0.454 |
| ALI x Spatial cue | -0.014 | 0.015 | -0.039 – 0.010 | -0.961 | 0.337 | 0.003 | 0.015 | -0.022 – 0.027 | 0.176 | 0.860 |
| ALI x Semantic cue | 0.027 | 0.015 | 0.003 – 0.051 | 1.838 | 0.066 | 0.019 | 0.015 | -0.005 – 0.043 | 1.287 | 0.198 |
| ALI x Probed ear | 0.009 | 0.015 | -0.016 – 0.034 | 0.570 | 0.569 | 0.002 | 0.015 | -0.023 – 0.027 | 0.113 | 0.910 |
| ALI x Age | 0.007 | 0.009 | -0.008 – 0.021 | 0.761 | 0.453 | -0.004 | 0.009 | -0.019 – 0.012 | -0.383 | 0.704 |
| **Random Effects** | | | | | | | | | | |
| $\sigma^2$ | 0.95 | | | | | 0.94 | | | | |
| $\tau_{00}$ | 0.04 $_{subj}$ | | | | | 0.06 $_{subj}$ | | | | |
| $\tau_{11}$ | 0.00 $_{subj.ALI\_norm\_zscored}$ | | | | | 0.00 $_{subj.ALI\_norm\_zscored}$ | | | | |
| $\rho_{01}$ | -0.09 $_{subj}$ | | | | | -0.48 $_{subj}$ | | | | |
| Observations | 17824 | | | | | 17824 | | | | |
| Marginal $R^2$ / Conditional $R^2$ | 0.005 / 0.051 | | | | | 0.003 / 0.065 | | | | |

#### Table S10: Brain-behavior model predicting accuracy including alpha power lateralization during cue presentation

| | Accuracy | | | | |
|---|---|---|---|---|---|
| *Predictors* | *Odds Ratios* | *std. Error* | *CI* | *z-value* | *p* |
| Intercept | 23.96 | 0.12 | 18.90 – 30.37 | 26.24 | **<0.001** |
| Spatial cue (selective) | 3.38 | 0.14 | 2.57 – 4.45 | 8.69 | **<0.001** |
| Semantic cue (specific) | 1.02 | 0.13 | 0.80 – 1.30 | 0.16 | 0.875 |
| Probed ear (right) | 1.38 | 0.09 | 1.15 – 1.66 | 3.51 | **<0.001** |
| First sentence (to-be-attended) | 1.00 | 0.06 | 0.88 – 1.13 | 0.01 | 0.995 |
| Age (z-scored) | 0.89 | 0.11 | 0.71 – 1.11 | -1.05 | 0.294 |
| PTA (z-scored) | 0.75 | 0.10 | 0.61 – 0.92 | -2.76 | **0.006** |
| Neural tracking attended (z-scored) | 0.99 | 0.03 | 0.92 – 1.06 | -0.41 | 0.679 |
| Neural tracking ignored (z-scored) | 1.04 | 0.03 | 0.97 – 1.11 | 1.05 | 0.294 |
| ALI cue (z-scored) | 1.00 | 0.03 | 0.93 – 1.06 | -0.15 | 0.882 |
| Probed ear x Semantic cue | 1.17 | 0.12 | 0.93 – 1.48 | 1.32 | 0.187 |
| First sentence x Semantic Cue | 2.17 | 0.12 | 1.71 – 2.75 | 6.41 | **<0.001** |
| Age x Spatial cue | 1.14 | 0.08 | 0.98 – 1.32 | 1.74 | 0.081 |
| Probed ear x Neural tracking attended | 1.18 | 0.07 | 1.03 – 1.35 | 2.37 | **0.018** |
| Probed ear x Neural tracking ignored | 0.85 | 0.07 | 0.74 – 0.97 | -2.36 | **0.018** |
| Probed ear x ALI cue | 0.94 | 0.07 | 0.83 – 1.07 | -0.90 | 0.366 |
| Neural tracking attended x ALI cue | 0.96 | 0.04 | 0.89 – 1.03 | -1.23 | 0.218 |
| Neural tracking ignored x ALI cue | 1.00 | 0.03 | 0.93 – 1.07 | -0.08 | 0.940 |
| Probed ear x Neural tracking attended x ALI cue | 0.98 | 0.07 | 0.85 – 1.13 | -0.28 | 0.779 |
| Probed ear x Neural tracking ignored x ALI cue | 0.97 | 0.07 | 0.85 – 1.11 | -0.40 | 0.688 |
| **Random Effects** | | | | | |
| $\sigma^2$ | 3.29 | | | | |
| $\tau_{00\ pair}$ | 0.66 | | | | |
| $\tau_{00\ subj}$ | 0.73 | | | | |
| $\tau_{11\ subj.spatialSelective}$ | 0.10 | | | | |
| $\tau_{11\ subj.probedRight}$ | 0.26 | | | | |
| $\rho_{01\ subj.spatialSelective}$ | 0.64 | | | | |
| $\rho_{01\ subj.probedRight}$ | 0.09 | | | | |
| Observations | 17532 | | | | |
| Marginal $R^2$ / Conditional $R^2$ | 0.108 / 0.385 | | | | |

**Table S11: Brain-behavior model predicting response speed including alpha power lateralization during cue presentation**

| | **Speed (z-scored)** | | | | |
|---|---|---|---|---|---|
| *Predictors* | *Estimates* | *std. Error* | *CI* | *t-value* | *p* |
| Intercept | -0.034 | 0.055 | -0.142 – 0.074 | -0.620 | 0.535 |
| Spatial cue (selective) | 0.577 | 0.050 | 0.479 – 0.674 | 11.595 | **<0.001** |
| Semantic cue (specific) | 0.192 | 0.035 | 0.122 – 0.261 | 5.413 | **<0.001** |
| Probed ear (right) | 0.063 | 0.015 | 0.033 – 0.093 | 4.123 | **<0.001** |
| First sentence (to-be-attended) | 0.008 | 0.013 | -0.017 – 0.033 | 0.609 | 0.542 |
| Age (z-scored) | -0.118 | 0.040 | -0.196 – -0.040 | -2.967 | **0.003** |
| PTA (z-scored) | -0.035 | 0.039 | -0.112 – 0.042 | -0.884 | 0.377 |
| Neural tracking attended (z-scored) | 0.003 | 0.007 | -0.011 – 0.016 | 0.400 | 0.689 |
| Neural tracking ignored (z-scored) | 0.001 | 0.007 | -0.012 – 0.015 | 0.207 | 0.836 |
| ALI cue (z-scored) | 0.003 | 0.006 | -0.009 – 0.016 | 0.535 | 0.593 |
| Spatial cue x Semantic cue | 0.103 | 0.068 | -0.031 – 0.236 | 1.509 | 0.131 |
| ALI cue x Neural tracking attended | -0.009 | 0.007 | -0.021 – 0.004 | -1.339 | 0.180 |
| ALI cue x Neural tracking ignored | 0.008 | 0.006 | -0.005 – 0.020 | 1.194 | 0.232 |
| Neural tracking attended x Spatial Cue | -0.037 | 0.013 | -0.063 – -0.011 | -2.747 | **0.006** |
| Neural tracking ignored x Spatial Cue | 0.006 | 0.013 | -0.020 – 0.033 | 0.481 | 0.630 |
| Neural tracking attended x First sentence | 0.024 | 0.014 | -0.002 – 0.051 | 1.781 | 0.075 |
| Neural tracking ignored x First sentence | -0.014 | 0.013 | -0.040 – 0.012 | -1.050 | 0.294 |
| **Random Effects** | | | | | |
| $\sigma^2$ | 0.60 | | | | |
| $\tau_{00\ pair}$ | 0.06 | | | | |
| $\tau_{00\ subj}$ | 0.21 | | | | |
| $\tau_{11\ subj.spatialSelective}$ | 0.10 | | | | |
| $\tau_{11\ subj.semanticSpecific}$ | 0.01 | | | | |
| $\tau_{11\ subj.probedRight}$ | 0.01 | | | | |
| $\rho_{01\ subj.spatialSelective}$ | 0.77 | | | | |
| $\rho_{01\ subj.semanticSpecific}$ | 0.30 | | | | |
| $\rho_{01\ subj.probedRight}$ | 0.07 | | | | |
| Observations | 16128 | | | | |
| Marginal $R^2$ / Conditional $R^2$ | 0.110 / 0.407 | | | | |

#### Methods and materials

##### Sentence materials and recordings

Speech stimuli consisted of 240 pairs of short German declarative sentences of fixed syntactic structure. All sentences were five words long. They always began with a first name, followed by sequence of a transitive verb, a temporal adverb, a case- and gender-ambiguous numeral and finally a plural noun (e.g., "Anna zeichnete gestern drei Stühle"; literal translation "Anna drew yesterday three chairs"). Each position could be filled with one of ten word alternatives (see supplemental information for the full list of used words). The number of syllables at each sentence position was held constant to control for overall sentence length. All sentence contexts (i.e., consisting of the first four words) were semantically non-predictive but but yielded plausible combinations with each of the 120 different sentence-final nouns. The task-relevant, sentence-final nouns belonged to two overarching general semantic categories: natural and man-made. In each of the two general categories there were ten specific subcategories (e.g., pets, fruits, vegetables in the natural, or instruments, furniture, tools in the man-made category) that each consisted of six highly representative members derived from a pre-experiment questionnaire study. Each noun was used four times across the final pool of sentence pairs, but always combined with different sentence contexts. From all possible permutations, we created 240 sentence pairs that differed at every word position.

A trained female speaker of standard German recorded the individual sentences in a sound-attenuated recording chamber (sampling rate, 44kHz). Root mean square (rms) intensity (–26 dB Full Scale, FS) was equalized across all individual sentences. When combining the sentence recordings per pair, we temporally aligned them by the onset of the two sentence-final nouns to ensure their simultaneous presentation. This, however led to slight differences in the onset on the individual sentences. Crucially, the range and average sentence onset difference was similar for trials in which the probed (to-be-attended) sentence began earlier and those in which the unprobed (to-be-ignored) sentence began earlier (probed first: range: 0–580 ms, 162.1 ms ±124.6; unprobed first: 0–560 ms, 180.6 ms ±127.2). Sentence presentation was masked by continuous speech-shaped noise at a signal-to-noise-ratio of 0 dB. Noise onset was presented with a 50 ms linear onset ramp and preceded sentence onset by 200 ms. Final speech stimuli had an average length of 2512 ms (range: 2183–2963 ms). All participants listened to the same 240 sentence pairs but in subject-specific randomized order. In addition, across participants we balanced the assignment of sentences to the right and left ear, respectively.
